# Retrospective whole-genome sequencing analysis distinguished PFGE and drug resistance matched retail meat and clinical *Salmonella* isolates

**DOI:** 10.1101/356857

**Authors:** Andrea B. Keefer, Lingzi Xiaoli, Nkuchia M. M’ikanatha, Kuan Yao, Maria Hoffmann, Edward G. Dudley

## Abstract

Non-typhoidal *Salmonella* are a leading cause of outbreak and sporadic-associated foodborne illnesses in the U.S. These infections have been associated with a range of foods, including retail meats. Traditionally, pulsed-field gel electrophoresis (PFGE) and antibiotic susceptibility testing (AST) have been used to facilitate public health investigations of *Salmonella* infections. However, whole-genome sequencing (WGS) has emerged as an alternative tool that can be routinely implemented. To assess its potential in enhancing integrated surveillance in Pennsylvania, WGS was used to directly compare the genetic characteristics of 7 retail meat and 43 clinical historic *Salmonella* isolates, subdivided into three subsets based on PFGE and AST results, to retrospectively resolve their genetic relatedness and identify antimicrobial resistance (AMR) determinants. Single nucleotide polymorphism (SNP) analyses revealed the retail meat isolates within *S*. Heidelberg, *S*. Typhimurium var. O5- subset 1, and *S*. Typhimurium var. O5- subset 2 were separated from each primary PFGE pattern-matched clinical isolate by 6-12, 41-96, and 21-81 SNPs, respectively. Fifteen resistance genes were identified across all isolates, including *fosA7*, a gene only recently found in a limited number of *Salmonella* and a ≥ 95% phenotype to genotype correlation was observed for all tested antimicrobials. Moreover, AMR was primarily plasmid-mediated in *S*. Heidelberg and *S*. Typhimurium var. O5- subset 2; whereas, AMR was chromosomally-carried in *S*. Typhimurium var. O5- subset 1. Similar plasmids were identified in both the retail meat and clinical isolates. Collectively, these data highlight the utility of WGS in retrospective analyses and enhancing integrated surveillance of *Salmonella* from multiple sources.

## Importance

Due to its enhanced resolution, whole-genome sequencing has emerged as a public health tool that can be utilized for pathogen monitoring, outbreak investigations, and surveillance for antimicrobial resistance. This study demonstrated that historical isolates that are indistinguishable by pulsed-field gel electrophoresis, a conventional genotyping method, and antibiotic susceptibility testing, could in fact be different strains, further highlighting the power of whole-genome sequencing. Moreover, we evaluated the role of whole-genome sequencing in integrated surveillance for drug-resistant *Salmonella* from retail meat and clinical sources in Pennsylvania and found a high correlation between antimicrobial resistance phenotype, as determined by antibiotic susceptibility testing, and genotype. Furthermore, the genomic context of each resistance gene was elucidated, which is critical to understanding how resistance is spreading within *Salmonella* in Pennsylvania. Taken together, these results demonstrate the utility and validity of whole-genome sequencing in characterizing human and food-derived *Salmonella*.

## Introduction

Non-typhoidal *Salmonella enterica* subsp. *enterica* are the leading bacterial etiological agent of foodborne illness, hospitalization, and death in the U.S. (1). The Centers for Disease Control and Prevention (CDC) estimates that non-typhoidal *Salmonella* cause 1.2 million infections, 23,000 hospitalizations, and 450 deaths annually (2). Furthermore, compared to other bacterial pathogens, non-typhoidal *Salmonella* account for the majority of foodborne outbreaks that occur in the U.S., with associated food commodities including eggs, vegetables, fruits, and retail meats (3, 4). Two important serovars of *Salmonella* are S. Typhimurium, including its variant, S. Typhimurium var. O5-, and S. Heidelberg; these serovars are consistently ranked within the top ten most commonly isolated from humans and retail meats in the U.S. (5, 6).

Although *Salmonella* infections are typically self-limiting, antimicrobial treatment can be necessary in some cases (7); accordingly, drug-resistant non-typhoidal *Salmonella* are categorized by the CDC as a serious public health threat (2). Indeed, an estimated 100,000 drug resistant non-typhoidal *Salmonella* infections occur annually in the U.S. (2). Specifically in Pennsylvania, to contribute to the One Health approach outlined by the White House for combating antimicrobial resistance (AMR), the Pennsylvania Department of Health (PADOH) conducts integrated AMR surveillance in enteric bacteria, including *Salmonella*, isolated from clinical samples and retail meats, as part of the National Antimicrobial Resistance Monitoring System (NARMS) (8, 9).

Moreover, the current standard methods employed to conduct integrated surveillance and foodborne outbreak investigations are antibiotic susceptibility testing (AST) and pulsed-field gel electrophoresis (PFGE). Even though PFGE has been considered the gold standard molecular epidemiological tool for decades, it has numerous documented limitations (10), including the inability to differentiate between clonal or low genetic diversity isolates, such as *S*. Heidelberg (11). Similarly, despite its utility, AST has several shortcomings, including MIC breakpoint inconsistencies (12), inability to efficiently test all known drugs, and only providing phenotype-level resolution (13), which is not adequate, if AMR gene alleles and/or transmission mechanisms need to be discerned.

Due to increases in affordability and ease of performance, whole-genome sequencing (WGS) has emerged as an attractive tool that can be utilized for foodborne outbreak investigations and pathogen-specific surveillance. Compared to conventional subtyping methods, WGS yields increased discriminatory power, which results from its nucleotide-level resolution, enabling the differentiation between clonal and closely related bacterial isolates (11, 14–17). Accordingly, prior studies have assessed and established the utility of WGS in *Salmonella* outbreak investigations, primarily by performing WGS retroactively on outbreak-associated isolates from various serovars (11, 14–22). WGS is also effective at source-tracking (20), as it provides the resolution needed to establish a genetic link between clinical and food isolates, which traditionally, can be difficult to attain (17, 18).

In addition, WGS has significant potential for use as an AMR surveillance tool that can be used to monitor and track resistance in human, animal, and food isolates. Indeed, NARMS has recently incorporated WGS into its AMR monitoring efforts (23). WGS allows for the elucidation of a bacterium’s full resistome; this information can be used to associate resistance phenotype with genotype and reveal possible transmission mechanisms, by characterizing the genomic context of each AMR gene (13, 24, 25).

Due to the enhanced resolution conferred by WGS, there is now motivation to re-examine historic collections of isolates to further resolve their relatedness and genetic AMR profiles.

Therefore, this study aimed to 1) use WGS to retrospectively resolve the genetic relatedness of a historic collection of PFGE-matched and multi-drug resistant (MDR) retail meat and clinical S. Heidelberg and S. Typhimurium var. O5- isolates, respectively and 2) to ascertain the genetic AMR profile of each isolate in that collection, in an effort to assess the role of WGS in integrated surveillance for drug-resistant *Salmonella* from clinical and retail meat sources in Pennsylvania.

## Materials and Methods

### Bacterial isolates

All bacterial isolates (n = 50) sequenced in this study, along with their associated metadata, are listed in Table 1. Henceforth, isolates SH-01 through SH-12 will collectively be referred to as the *S.* Heidelberg subset; isolates SC-01 through SC-28 will be referred to as *S.* Typhimurium var. O5- subset 1; and isolates SC-29 through SC-38 will be referred to as *S.* Typhimurium var. O5- subset 2.

**Table 1.**
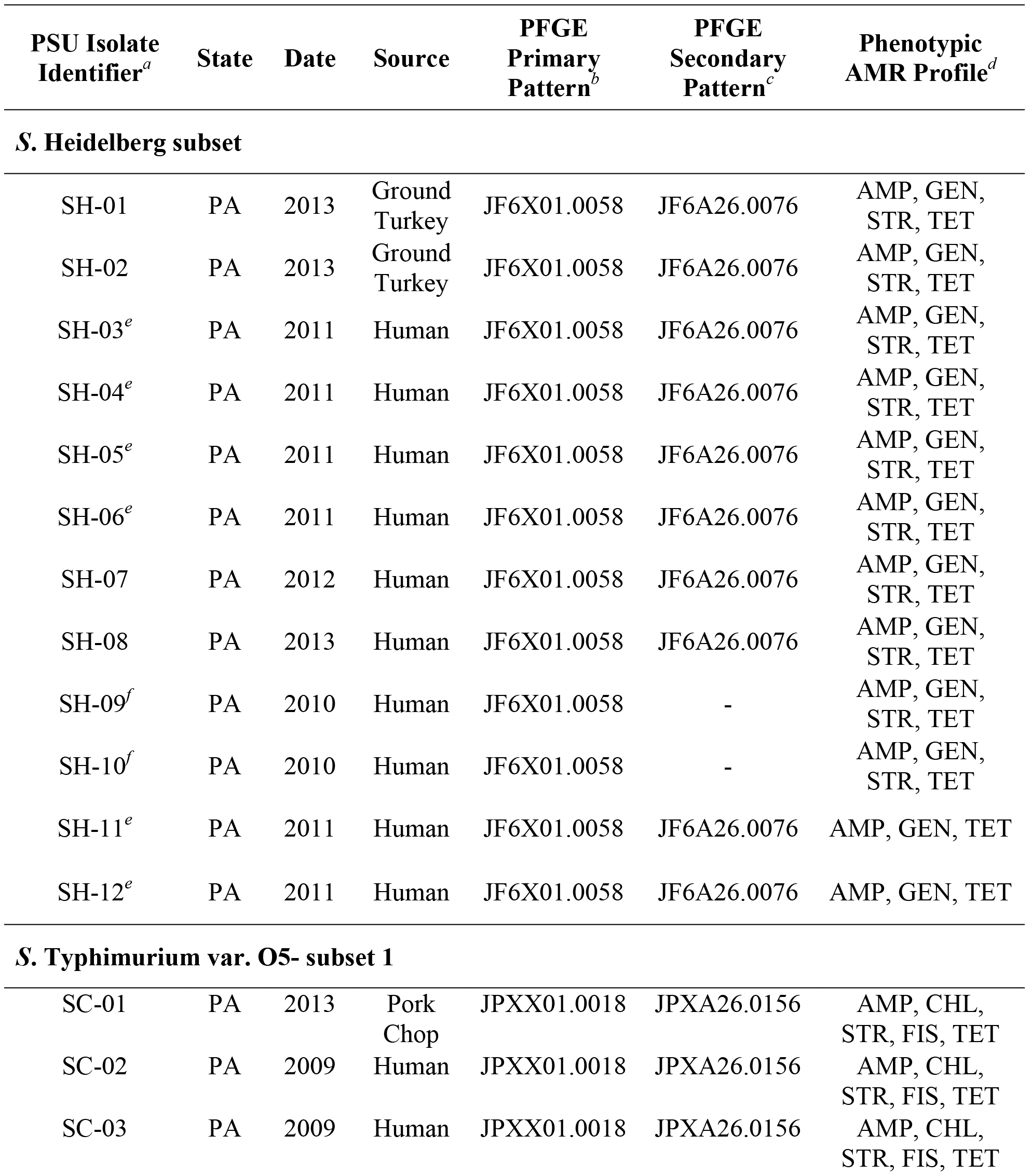
Metadata of *S*. Heidelberg and *S*. Typhimurium var. O5- isolates sequenced in this study.

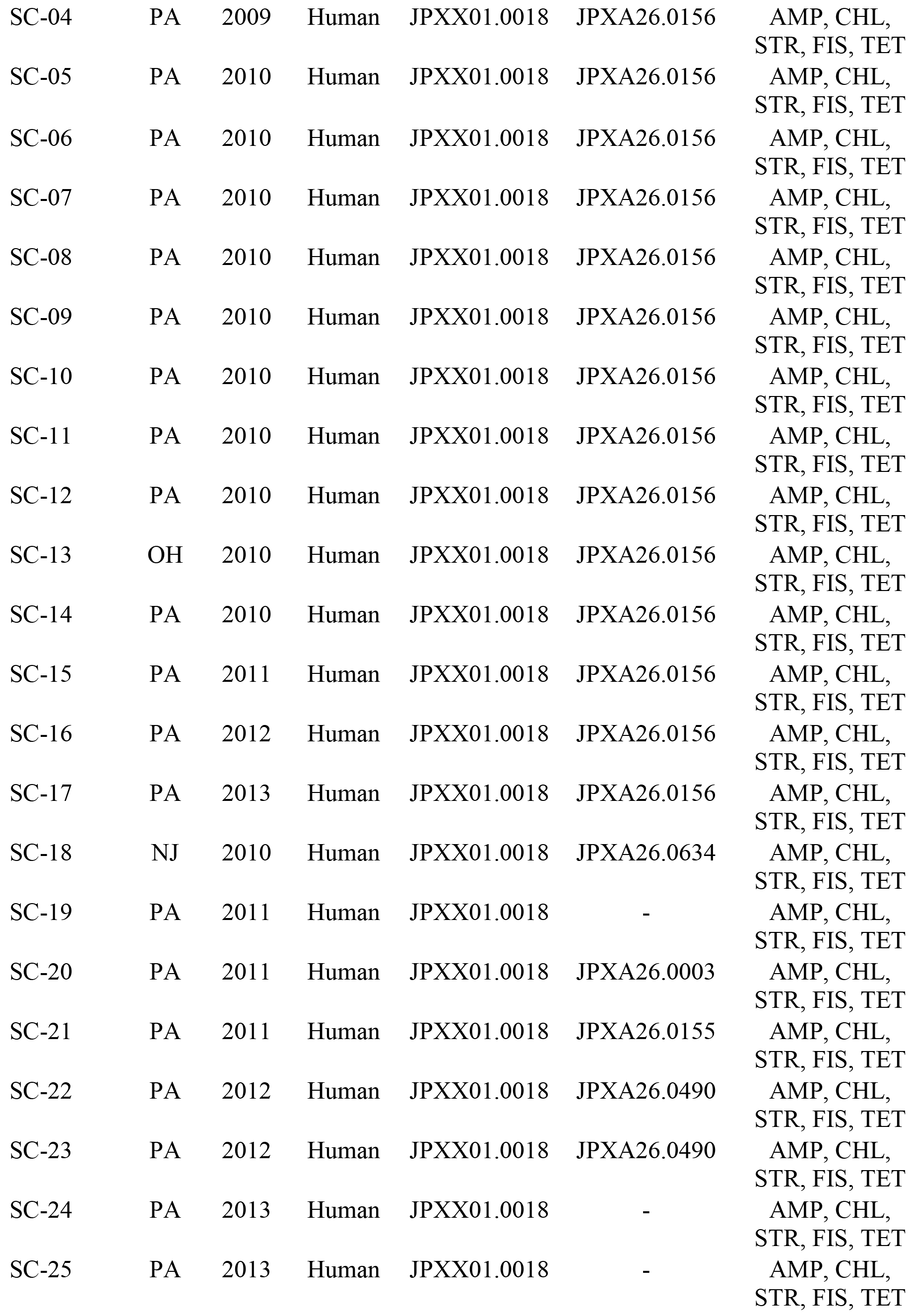

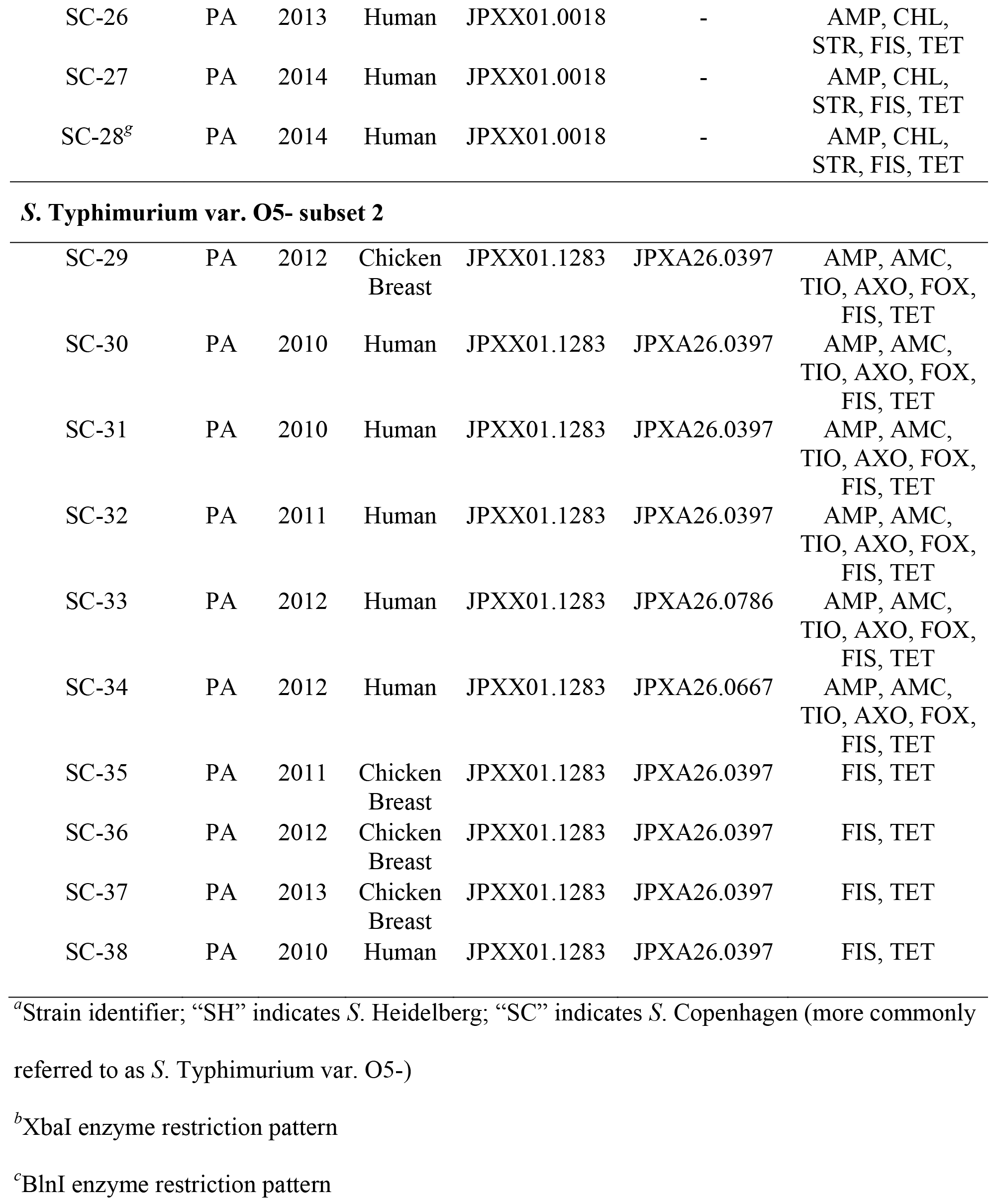

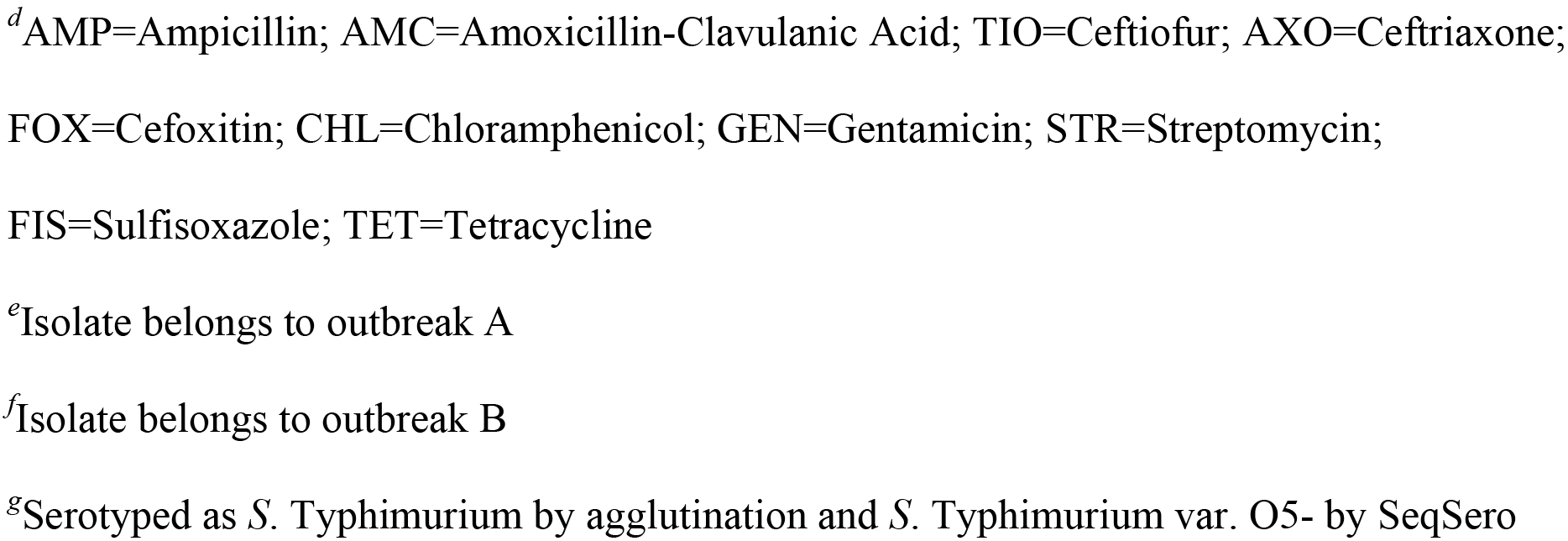

As part of NARMS surveillance, isolates SH-01, SH-02, SC-01, SC-29, and SC-35 through SC-37 were recovered and identified from retail meats (ground turkey, pork chop, and chicken breast), purchased throughout the state of Pennsylvania between 2009-2014, following standard protocols by the PADOH (26). Of note, the S. Heidelberg retail meat isolates were derived from meats processed at different facilities; however, the S. Typhimurium var. O5-subset 2 retail meat isolates SC-29, SC-35, and SC-36 were all derived from meats that were originally processed at the same facility. All clinical *Salmonella* were from a collection of human isolates that had indistinguishable PFGE patterns with retail meat isolates collected as part of NARMS in Pennsylvania; the clinical isolates were submitted to the PADOH Bureau of Laboratories by clinical laboratories in compliance with public health reporting requirements (27).

### Pulsed-field gel electrophoresis (PFGE) and antibiotic susceptibility testing (AST)

All retail meat and clinical isolates previously underwent PFGE and AST characterization through the PADOH. Each retail meat and clinical isolate was subjected to PFGE using the XbaI restriction enzyme, following the CDC’s PulseNet protocol for *Salmonella* subtyping (28). For all isolates, except SH-09, SH-10, SC-19, and SC-24 through SC-28, the BlnI restriction enzyme was also used to obtain a secondary PFGE pattern. All retail meat and matched clinical isolates were also subjected to AST as previously described (29) for susceptibility to the antimicrobial agents: gentamicin, streptomycin, ampicillin, amoxicillin-clavulanic acid, cefoxitin, ceftiofur, ceftriaxone, azithromycin, chloramphenicol, nalidixic acid, ciprofloxacin, sulfisoxazole, trimethoprim-sulfamethoxazole, and tetracycline. Resistance was defined using CLSI criteria if available, otherwise NARMS interpretative criteria were used (30).

### Genomic DNA extraction, library construction, and genome sequencing

A 3 mL Lysogeny Broth (LB) overnight culture, derived from a single colony, was prepared for each isolate. Total genomic DNA was extracted using the Wizard© Genomic DNA Purification Kit (Promega, Madison, WI, USA) following the manufacturer’s instructions. Genomic DNA purity was confirmed via an A_260_/A_280_ measurement (target ≥ 1.8) and the concentration was determined using the Qubit™ dsDNA Broad-Range quantification kit (Thermo Fisher Scientific, Waltham, MA, USA). Following quantification, genomic DNA was diluted to 0.2 ng/μL. A paired-end DNA library was prepared and normalized using the Nextera XT DNA Library Prep Kit (Illumina, Inc., San Diego, CA, USA). The resulting library was sequenced on an Illumina MiSeq sequencer (Illumina, Inc., San Diego, CA, USA) using a MiSeq reagent v2, 500-cycle kit, with 250 bp read length. Additionally, a representative isolate from each subset (SH-04, SC-09, and SC-31, respectively) was sequenced on a PacBio *RS* II (Pacific Biosciences, Menlo Park, CA, USA) as previously described (31). Specifically, we prepared the library using 10 μg genomic DNA that was sheared to a size of 20-kb fragments by g-tubes (Covaris, Inc., Woburn, MA, USA) according to the manufacturer’s instruction. The SMRTbell 20-kb template library was constructed using DNA Template Prep Kit 1.0 with the 20-kb insert library protocol (Pacific Biosciences, Menlo Park, CA, USA). Size selection was performed with BluePippin (Sage Science, Beverly, MA, USA). The library was sequenced using the P6/C4 chemistry on 2 single-molecule real-time (SMRT) cells with a 240-min collection protocol along with stage start. Analysis of the sequence reads was implemented using SMRT Analysis 2.3.0. The best *de novo* assembly was established with the PacBio Hierarchical Genome Assembly Process (HGAP3.0) program, which resulted in the closed chromosome of each isolate (Table S1). Each closed chromosome was annotated using Rapid Annotation using Subsystem Technology (RAST) (32).

### Sequencing quality control

Following sequencing, Illumina read quality was confirmed using FastQC v0.11.5 (33). Raw Illumina reads for each isolate within each subset were aligned to the closed chromosome of SH-04, SC-09, and SC-31, respectively using Burrows-Wheeler Aligner v0.7.15 (BWA-MEM) (34). Subsequently, average genome coverage was calculated using the SAMtools v1.4 depth command (35). Additionally, Illumina reads from each isolate were *de novo* assembled using SPAdes v3.9 (36). QUAST v4.5 (37) was used to assess assembled/draft genome quality. All isolates sequenced in this study by Illumina technology had > 30X coverage, < 200 contigs, an N50 score > 200,000, and total assembly length between 4.4-5.l megabases (Mb) (Fig. S1A, B). The previously determined agglutination derived serotype of each isolate was also confirmed using SeqSero (38).

### Comparative SNP and phylogenetic analyses

The validated single nucleotide polymorphism/variant (SNP or SNV) calling pipeline SNVPhyl v1.0.1 (39) was utilized to identify variants. Default parameters were used with the exception of the following: minimum coverage was set to 10X, minimum mean mapping was set to 30, and the SNV abundance ratio was set to 0.75. Preliminarily, the assembled genomes (from Illumina sequencing) of SH-04, SC-09, and SC-38 were used as the reference genomes for SNP and phylogenetic analyses in each subset (data not shown). Subsequently, the complete chromosome sequences (from PacBio sequencing) of isolates SH-04, SC-09, and SC-31 were used as the reference genomes for the *S.* Heidelberg subset, *S.* Typhimurium var. O5- subset 1, and *S.* Typhimurium var. O5- subset 2, respectively. The SNVPhyl pipeline outputted a concatenated SNP alignment file, a pairwise SNP distance matrix, and a maximum likelihood phylogenetic tree generated using PhyML v3.1.1 for each subset. The SNP alignment file was also used to construct a custom maximum likelihood phylogenetic tree using PhyML v3.1.1 (40). This tree was constructed using the GTR + gamma model with 1,000 bootstrap replicates.

### SNP annotation

Concatenated base call files from SNVPhyl were downloaded and converted to VCF files using BCFtools v1.3.1 view (35); vcf-subset v0.1.13 (41) was used to filter out all non-variant positions. The filtered VCF files for each isolate were combined into a single VCF file using BCFtools merge for each subset. A custom SnpEff database was built for annotation of each subset using the RAST-generated GenBank file for either SH-04, SC-09, or SC-31. SNPs were then annotated using SnpEff v4.3 (42). SNP annotations were subsequently filtered to only include valid SNPs, as determined by SNVPhyl. Gene names for each valid SNP were extracted from the RAST-generated GFF file of either SH-04, SC-09, or SC-31. SNP annotation script can be found at https://github.com/DudleyLabPSU/SNP-Annotation.

### Genetic AMR profile determination

Genetic AMR determinants were identified in all genomes, using the Bacterial Antimicrobial Resistance Reference Gene Database (BARRGD) (Accession number PRJNA313047; accessed April 2018) and BLAST+ (43). To confirm BARRGD results and determine if any AMR-associated chromosomal point mutations were present, ResFinder 3.0 (44) was used. AMR genes that were detected by either method were only considered to be present if they had ≥ 90% nucleotide identity and ≥ 60% coverage (these parameters align with ResFinder’s default search settings).

### Plasmid identification and characterization

Known plasmid replicon sequences were identified using the PlasmidFinder (45) database and BLAST+. BLAST alignments were filtered to only include those present at ≥ 95% nucleotide identity and ≥ 60% coverage (these parameters align with PlasmidFinder’s default search settings), with the exception of IncX1 in the S. Heidelberg subset, which was included due to its presence at 94.9% nucleotide identity. PacBio sequencing and subsequent processing (as described above) were also used to attain complete plasmid sequences within SH-04, SC-09, and SC-31 (Table S1). Each closed plasmid sequence was annotated using RAST (32). BLAST Ring Image Generator (BRIG) (46) was used to visualize plasmid comparisons in all subsets. Complete plasmid sequences were also compared to plasmids deposited in NCBI’s GenBank database (Table S2) using BLAST (47) and BRIG, to assess novelty.

### Accession numbers

Raw Illumina WGS data were submitted to NCBI and subsequently, assigned BioSample and SRA accession numbers (Table S3); all data can also be found under BioProject PRJNA357723. The closed chromosome and plasmid sequences of isolates SH-04, SC-09, and SC-31 were submitted to GenBank and their accession numbers are also found in Table S3.

## Results

### Comparison of retail meat and human isolates based on PFGE and AST results

In total, the PFGE patterns of 86 retail meat isolates were indistinguishable from the PFGE patterns of 1,665 clinical isolates located in the Pennsylvania surveillance database. From that larger collection, three subsets were chosen to whole-genome sequence and are referred to as the S. Heidelberg subset, S. Typhimurium var. O5- subset 1, and S. Typhimurium var. O5- subset 2. These subsets were selected based on quantity (≥ one retail meat matching multiple clinical isolates), AMR (primarily identical MDR patterns within each subset), and serovar relevance.

All isolates within the *S*. Heidelberg subset (two retail meat and ten clinical) matched by primary PFGE pattern (JF6X01.0058) and all isolates, except SH-09 and SH-10 (secondary patterns unknown), shared the same secondary PFGE pattern (JF6A26.0076) (Table 1). Isolates SH-01 through SH-10 displayed resistance to ampicillin, gentamicin, streptomycin, and tetracycline antimicrobials; whereas, isolates SH-11 and SH-12 were only resistant to ampicillin, gentamicin, and tetracycline (Table 1). All *S.* Typhimurium var. O5- subset 1 isolates (one retail meat and 27 clinical) shared the same primary PFGE pattern (JPXX01.0018) and isolates SC-01 through SC-17 also shared the same secondary PFGE pattern (JPXA26.0156) (Table 1). Additionally, all isolates within this subset were phenotypically resistant to ampicillin, chloramphenicol, streptomycin, sulfisoxazole, and tetracycline (ACSSuT pattern) (Table 1). All *S.* Typhimurium var. O5- subset 2 isolates (four retail meat and six clinical) matched by primary PFGE pattern (JPXX01.1283) and all isolates except SC-33 and SC-34 matched by secondary PFGE pattern (JPXA26.0397) (Table 1). However, this subset was defined by two distinct AMR profiles: isolates SC-29 through SC-34 displayed resistance to ampicillin, amoxicillin-clavulanic acid, ceftiofur, ceftriaxone, cefoxitin, sulfisoxazole, and tetracycline; whereas, isolates SC-35 through SC-38 were resistant to only sulfisoxazole and tetracycline antimicrobials (Table 1).

### Comparative SNP analysis resolved phylogenetic relationships in each subset

To retrospectively resolve the genetic relatedness of each PFGE-matched retail meat and clinical isolate within all three subsets, the validated SNP-calling pipeline, SNVPhyl was utilized. In total, the S. Heidelberg subset was defined by 35 SNPs. These SNPs were generally distributed over the length of the reference chromosome (Fig. S2A, top) and were predominantly non-synonymous (Fig. S2B); notably, SNP annotation revealed that missense mutations were located in flagellar-associated genes (*fliC* and *fliE*) in some isolates. In terms of SNP distances, the retail meat isolates (SH-01 and SH-02) were separated from their ten PFGE-matched clinical isolates by 6 to 12 SNPs (Table 2). Interestingly, despite being derived from two different processing facilities, the two retail meat isolates were separated from one another by 1 unique SNP (Table 2); a missense variant (Ala138Thr) in *rapA*. Furthermore, within this subset, six clinical isolates were previously determined to be outbreak-associated (outbreak A). These isolates (SH-03 through SH-06, SH-11, and SH-12) were separated from each other by 3 to 9 SNPs (Table 2). Additionally, clinical isolates SH-09 and SH-10 were previously determined to be part of a separate outbreak (outbreak B). Indeed, under these experimental conditions, they were separated by 0 SNPs (Table 2). Moreover, to visualize the phylogenetic relationships of the *S*. Heidelberg isolates, a maximum likelihood phylogenetic tree was constructed. The retail meat isolates branched distinctly away from the *S*. Heidelberg clinical isolates (Fig. 1). The clinical isolates all clustered on the same primary branch; notably, sporadic clinical isolate SH-08 clustered with the confirmed outbreak isolates (Fig. 1). Furthermore, the use of the draft genome of SH-04 as the reference genome resulted in nearly identical SNP distances and phylogenetic tree topologies; these similarities were also observed in each *S*. Typhimurium var. O5- subset, when the SC-09 or SC-38 assembled genome was utilized as the reference (data not shown).

**Figure 1.**
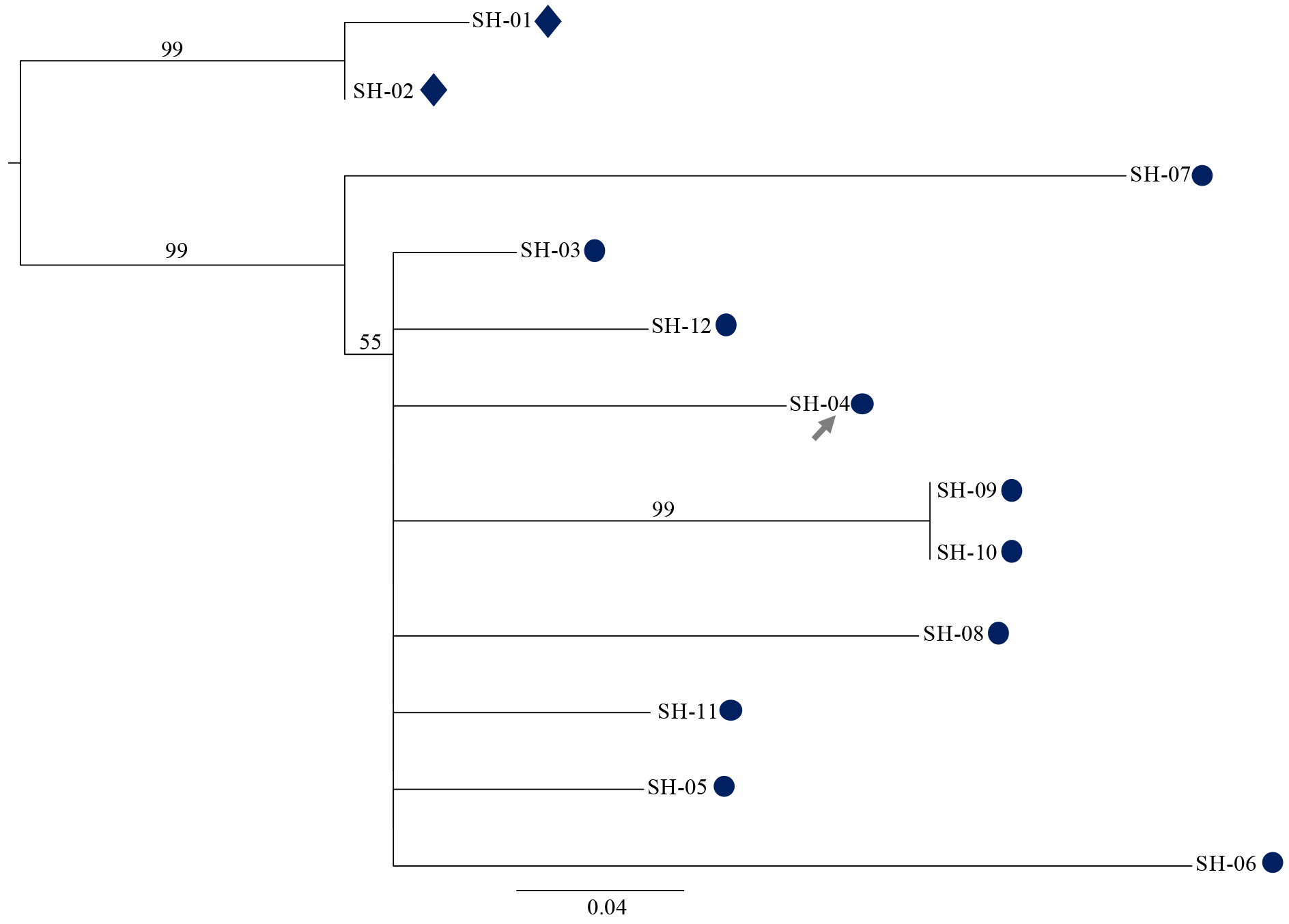
Phylogenetic relationships of *S*. Heidelberg isolates. Maximum likelihood phylogenetic tree of *S*. Heidelberg retail meat (diamonds) and clinical (circles) isolates generated by PhyML v3.1.1 (40) using the GTR + gamma model and 1,000 bootstrap replicates. Isolates SH-03 through SH-06, SH-11, and SH-12 were previously determined to be part of an outbreak (outbreak A); isolates SH-09 and SH-10 were previously determined to be part of a separate outbreak (outbreak B). The SNP alignment file produced by SNVPhyl v1.0.1 (39) contained 35 SNPs and served as the input for PhyML. The closed chromosome of clinical isolate SH-04 was used as the reference genome (grey arrow). Bootstrap values above 50 are included on the tree.

**Table 2.** Pairwise SNP distances between S. Heidelberg isolates.

|  | Retail Meat |  | Outbreak A Clinical |  |  |  | Clinical |  | Outbreak B Clinical |  | Outbreak A Clinical |  |
| --- | --- | --- | --- | --- | --- | --- | --- | --- | --- | --- | --- | --- |
|  | SH-01 | SH-02 | SH-03 | SH-04 | SH-05 | SH-06 | SH-07 | SH-08 | SH-09 | SH-10 | SH-11 | SH-12 |
| SH-01 | 0 |  |  |  |  |  |  |  |  |  |  |  |
| SH-02 | 1 | 0 |  |  |  |  |  |  |  |  |  |  |
| SH-03 | 7 | 6 | 0 |  |  |  |  |  |  |  |  |  |
| SH-04 | 9 | 8 | 4 | 0 |  |  |  |  |  |  |  |  |
| SH-05 | 8 | 7 | 3 | 5 | 0 |  |  |  |  |  |  |  |
| SH-06 | 12 | 11 | 7 | 9 | 8 | 0 |  |  |  |  |  |  |
| SH-07 | 10 | 9 | 7 | 9 | 8 | 12 | 0 |  |  |  |  |  |

| SH-08 | 10 | 9 | 5 | 7 | 6 | 10 | 10 | 0 |  |  |  |  |
| --- | --- | --- | --- | --- | --- | --- | --- | --- | --- | --- | --- | --- |
| SH-09 | 10 | 9 | 5 | 7 | 6 | 10 | 10 | 8 | 0 |  |  |  |
| SH-10 | 10 | 9 | 5 | 7 | 6 | 10 | 10 | 8 | 0 | 0 |  |  |
| SH-11 | 8 | 7 | 3 | 5 | 4 | 8 | 8 | 6 | 6 | 6 | 0 |  |
| SH-12 | 8 | 7 | 3 | 5 | 4 | 8 | 8 | 6 | 6 | 6 | 4 | 0 |

Furthermore, *S*. Typhimurium var. O5- subset 1 was defined by 482 SNPs that were uniformly distributed across the reference chromosome (Fig. S2A, middle). The majority of the SNPs were non-synonymous (Fig. S2C). Moreover, in some *S*. Typhimurium var. O5- subset 1 isolates, missense mutations were located in multiple fimbrial-associated genes (*fimD*, *stdB, fimF*), type III secretion system 1 (T3SS1)-associated genes (*spaN*, *sirC*), and flagellar-associated genes (*fliC*, *motA*). The overall pairwise SNP distances within this collection ranged from 0 to 105 (Table 3). Even though the retail meat isolate (SC-01) shared the same primary and secondary PFGE patterns and drug resistance profile as clinical isolates SC-02 through SC-17, on the genome level, it was surprisingly separated from each of those isolates by 41 to 75 SNPs (Table 3). Furthermore, SC-01 was separated from clinical isolates SC-18 through SC-28 by 46 to 96 SNPs, despite matching by primary PFGE pattern and drug resistance (Table 3). Nonetheless, some clinical isolates were genetically indistinguishable from one another (SC-05 and SC-06 and SC-14 and SC-19) or otherwise very closely related (shown in light grey shading in Table 3). A maximum likelihood tree was also constructed for visualization of phylogenetic relationships. Retail meat isolate SC-01 generally clustered with clinical isolates SC-02 through SC-06 and SC-17, but importantly, SC-01 was still separated from those isolates by 41 to 44 SNPs (Fig. 2 and Table 3). Interestingly, these isolates all originated from the southern portion of Pennsylvania. SC-01 and SC-06 were classified as southeast; SC-04 and SC-05 were classified as southcentral; SC-02, SC-03, and SC-17 were classified as southwest. The remainder of the clinical isolates branched separately, with some forming distinct clusters consistent with the pairwise SNP distances (Fig. 2).

**Figure 2.**
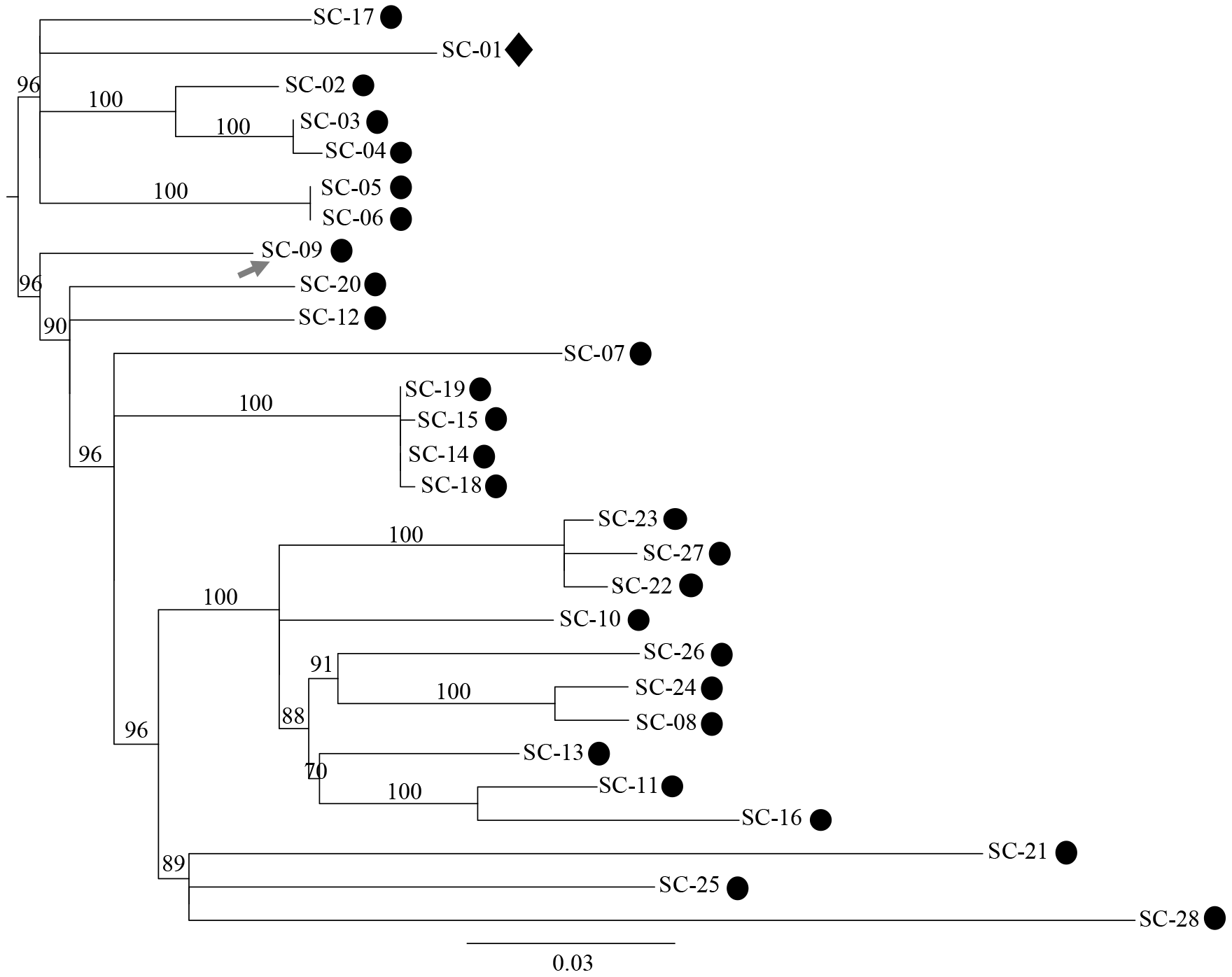
Phylogenetic relationships of *S*. Typhimurium var. O5- subset 1 isolates. Maximum likelihood phylogenetic tree of *S*. Typhimurium var. O5- subset 1 retail meat (diamond) and clinical (circles) isolates constructed by PhyML v3.1.1 (40) using the GTR + gamma model and 1,000 bootstrap replicates. The SNP alignment file produced by SNVPhyl v1.0.1 (39) served as the input for PhyML and contained 482 SNPs. The closed chromosome of clinical isolate SC-09 was used as the reference genome (grey arrow). Bootstrap values above 65 are included on the tree.

**Table 3.**
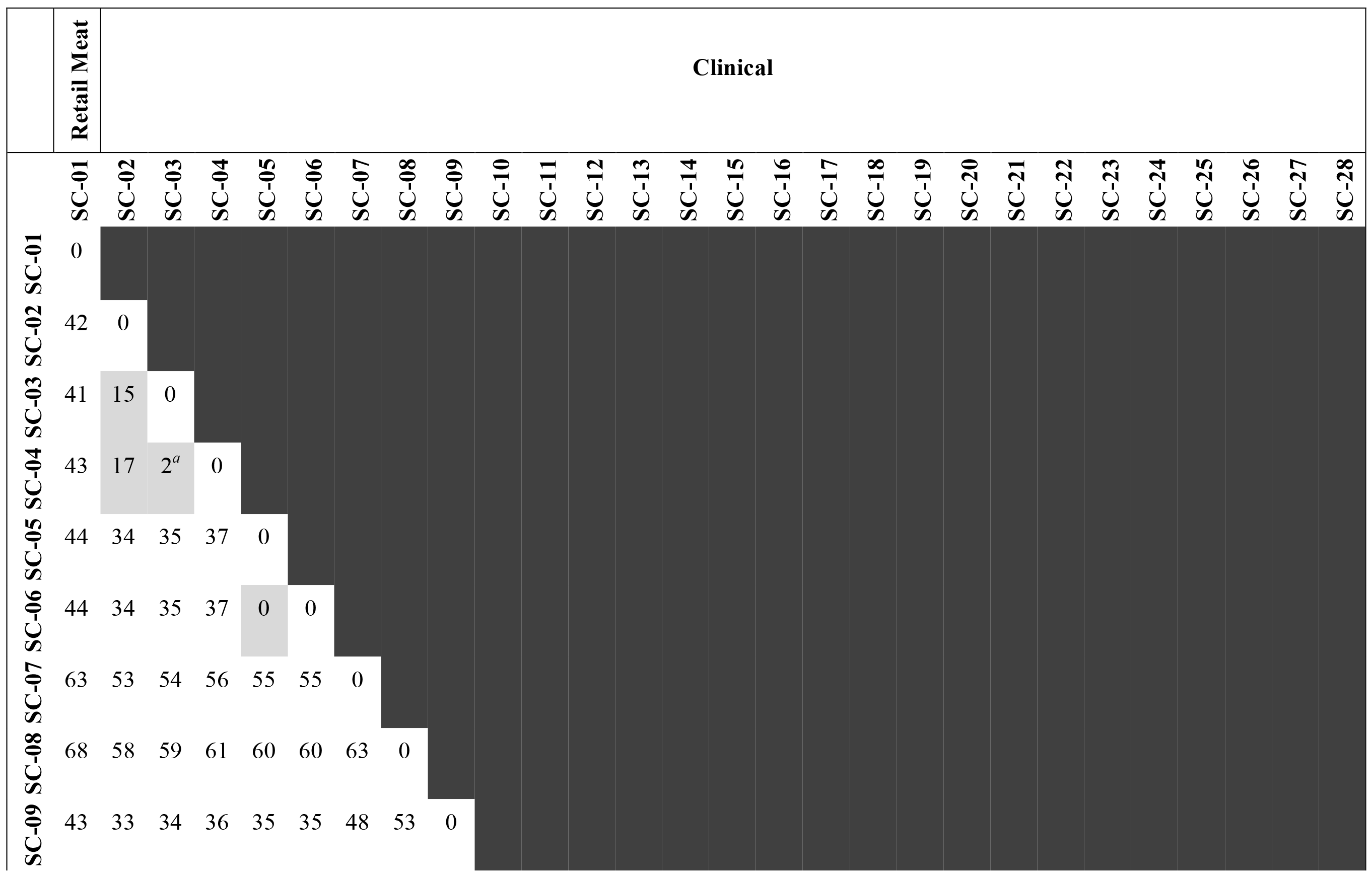
Pairwise SNP distances between S. Typhimurium var. O5- subset 1 isolates.

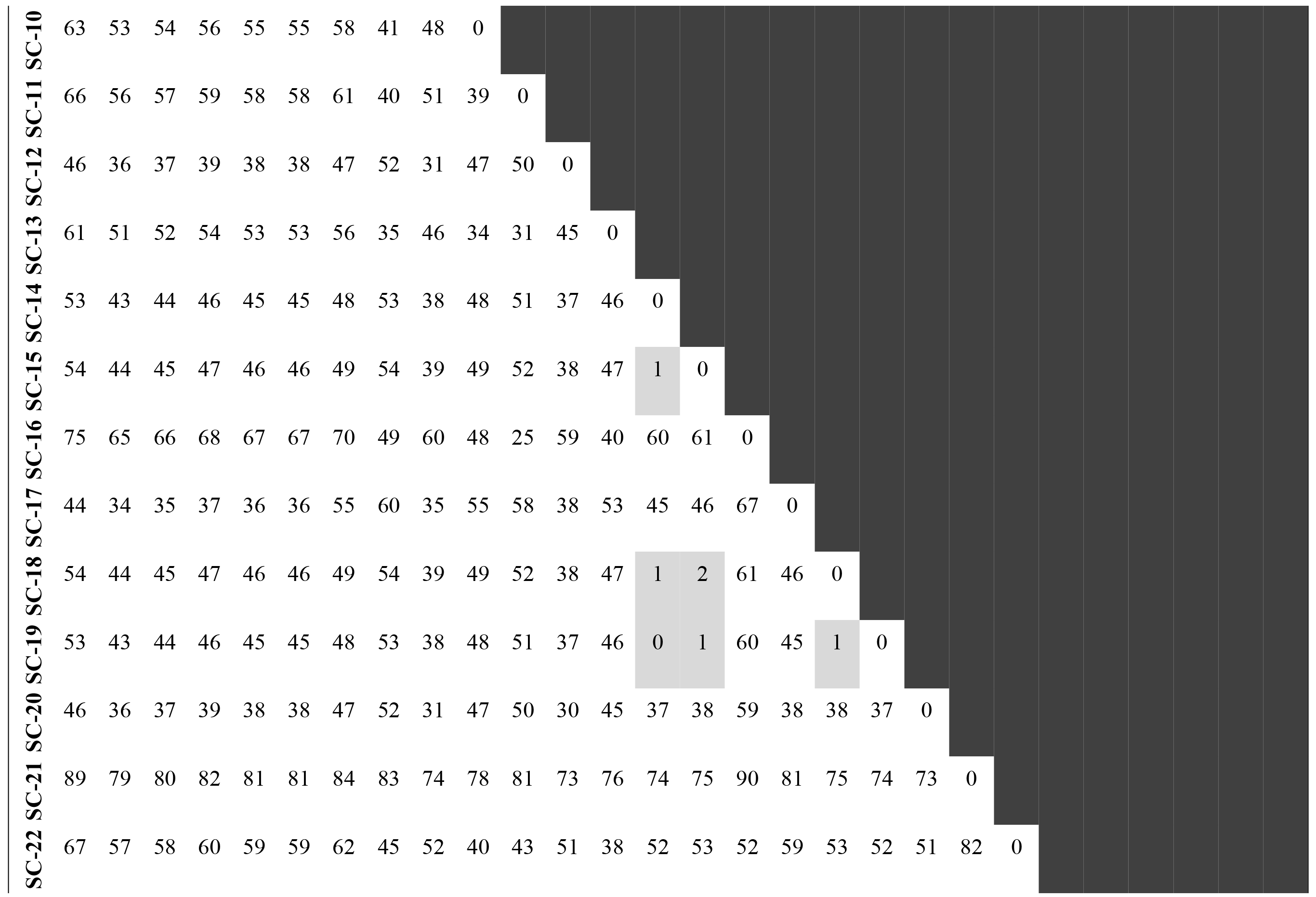

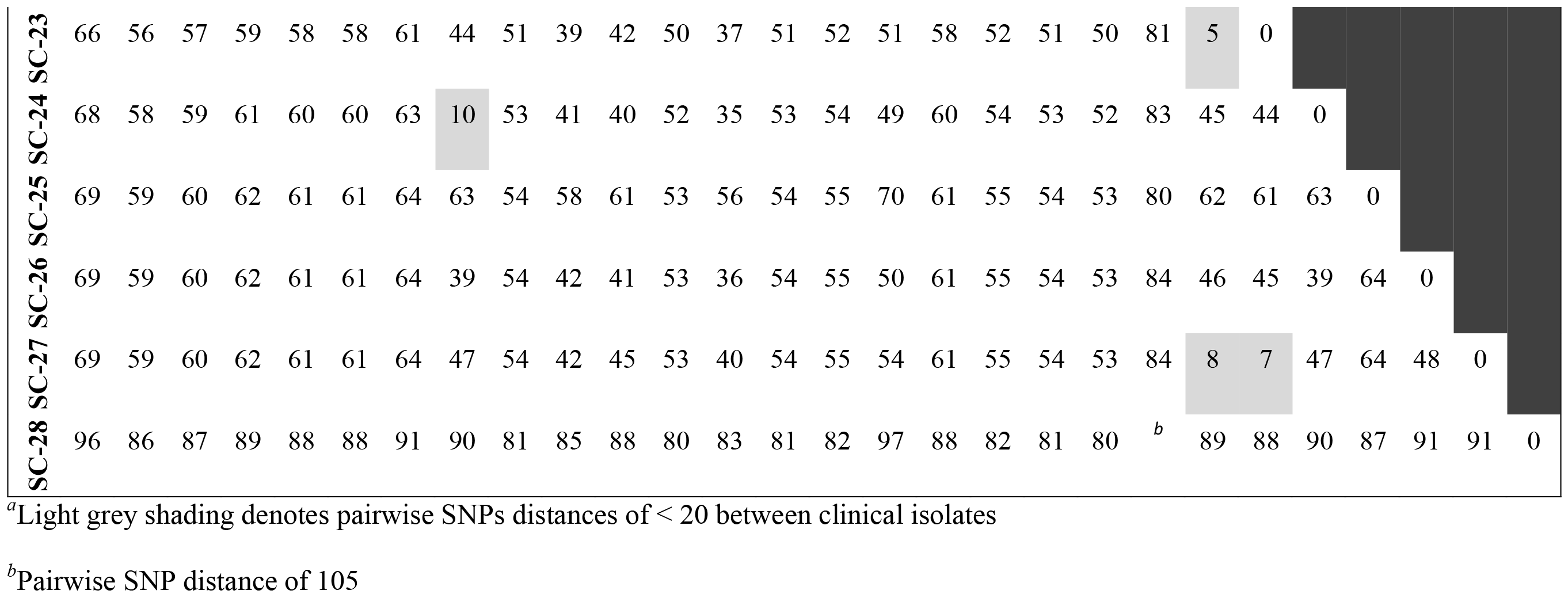

Lastly, *S*. Typhimurium var. O5- subset 2 was defined by 225 SNPs that were evenly dispersed across the reference chromosome (Fig. S2A, bottom). Similar to the previous two subsets, the majority of the SNPs were non-synonymous (Fig. S2D). Interestingly, in some *S*. Typhimurium var. O5- subset 2 isolates missense or nonsense mutations were located in genes that encode the T3SS1 effector *sipA* and the type III secretion system 2 (T3SS2) structural component and effector protein, *ssaV* and *sseF*. In terms of relatedness, the first retail meat isolate, SC-29, was separated from its primary PFGE, secondary PFGE, and AST-matched clinical isolates (SC-30 through SC-32) by 58, 61, and 27 SNPs, respectively (Table 4). The remaining three retail meat isolates, SC-35, SC-36, and SC-37, and clinical isolate SC-38 matched by both PFGE patterns and drug resistance. However, SC-38 was separated from SC-35, SC-36, and SC-37 by 24, 27, and 31 SNPs, respectively (Table 4). Of note, the smallest pairwise SNP distance between a clinical and retail meat isolate was 21, which occurred between clinical isolate SC-32 and retail meat isolate SC-35 (Table 4); these isolates matched by both PFGE patterns, but did not share the same resistance phenotype. In total, all four retail meat isolates were separated from the six clinical isolates in this subset by 21 to 81 SNPs (Table 4). When visualized on a phylogenetic tree, two distinct clusters of isolates were observed (Fig. 3). Clinical isolates SC-30, SC-31, SC-33, and SC-34 all distinctly branched away from the four retail meat isolates and furthermore, were fairly dissimilar from one another as well (Fig. 3). Conversely, clinical isolates SC-32 and SC-38 clustered with all four retail meat isolates, but importantly, these isolates were still separated from the retail meat isolates by 21 to 31 SNPs (Fig. 3 and Table 4). In general, the trend of relatedness was the same in both S. Typhimurium var. O5- subsets, in that, isolates that matched by conventional methods, were all separated by a relatively large number of SNPs, especially when compared to the S. Heidelberg collection.

**Figure 3.**
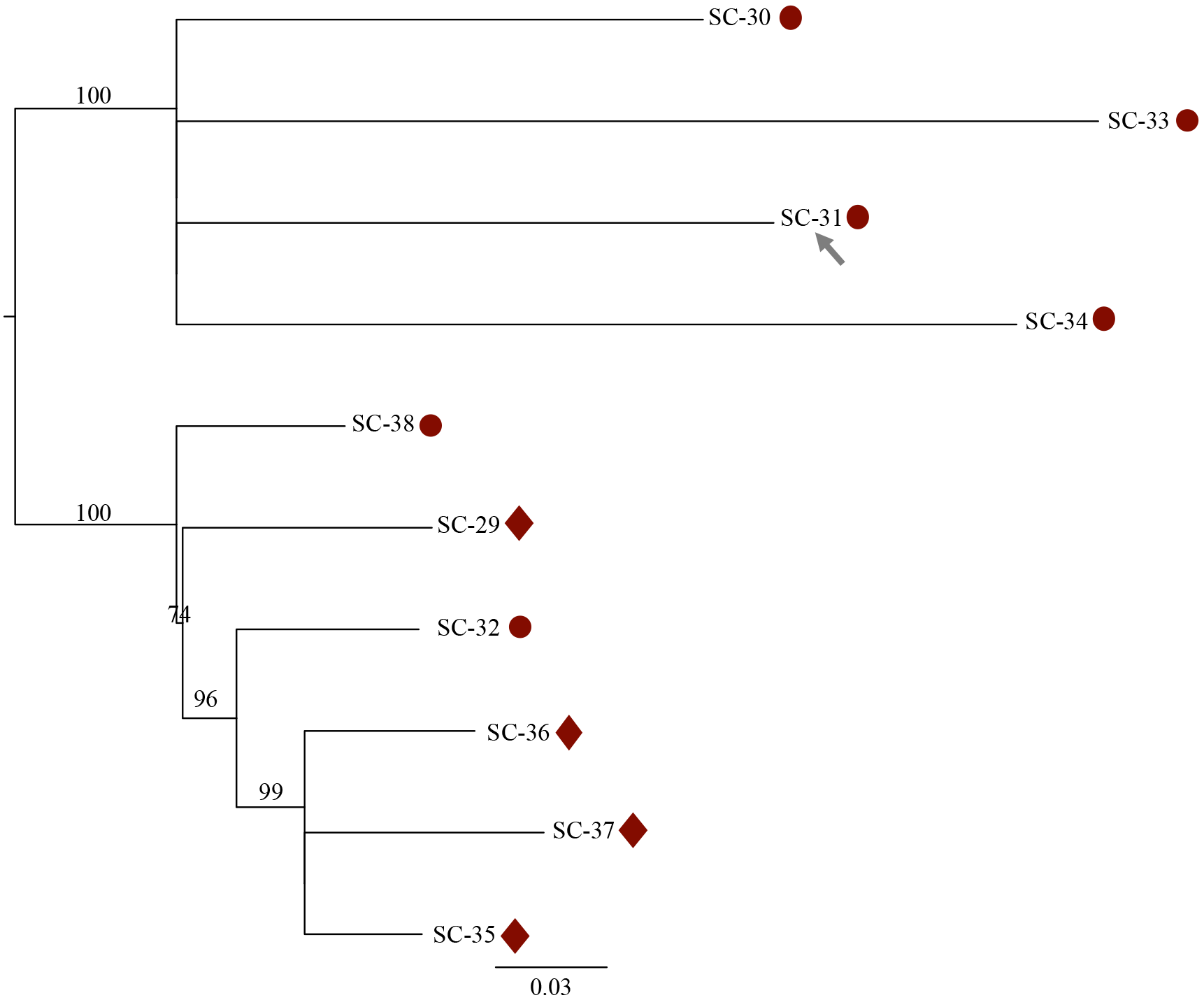
Phylogenetic relationships of *S*. Typhimurium var. O5- subset 2 isolates. Maximum likelihood phylogenetic tree of *S*. Typhimurium var. O5- subset 2 retail meat (diamonds) and clinical (circles) isolates constructed by PhyML v3.1.1 (40) using the GTR + gamma model and 1,000 bootstrap replicates. The SNP alignment file produced by SNVPhyl v1.0.1 (39) served as the input for PhyML and contained 225 SNPs. The closed chromosome of clinical isolate SC-31 was used as the reference genome (grey arrow). Bootstrap values above 70 are included on the tree.

**Table 4.**
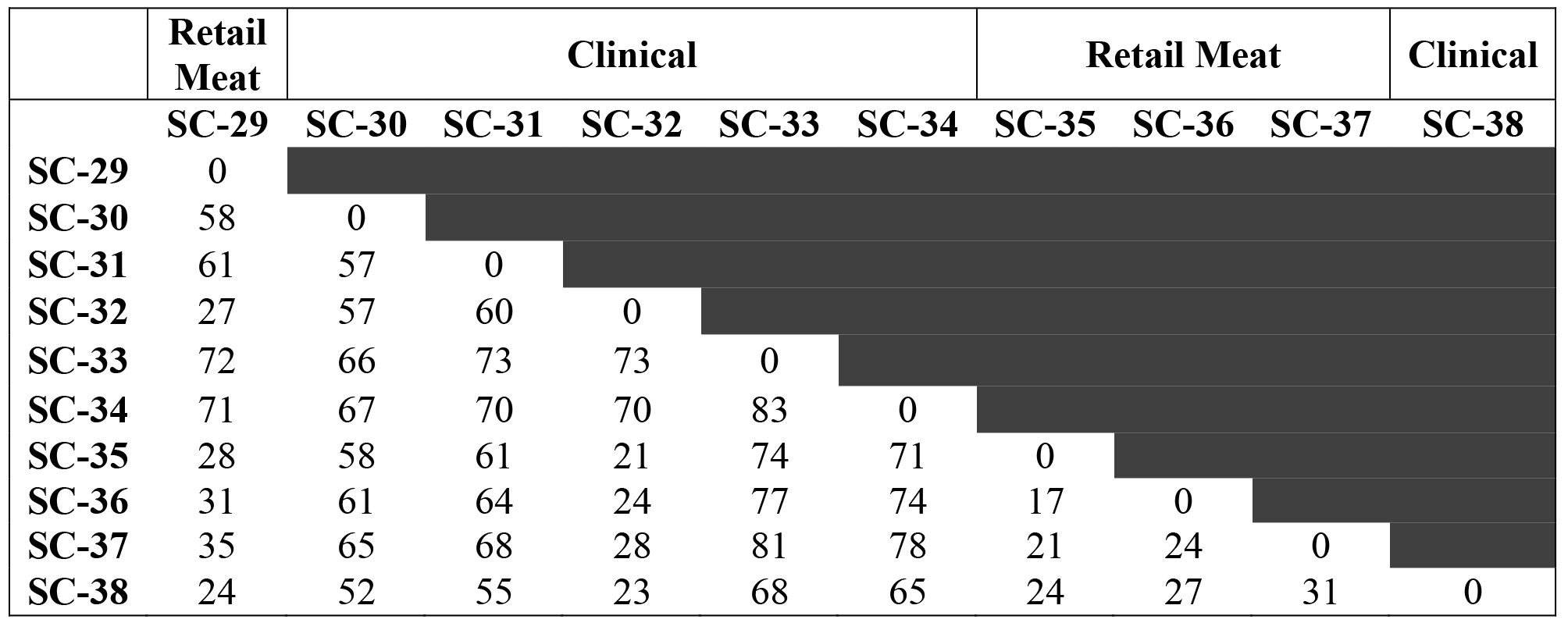
Pairwise SNP distances between S. Typhimurium var. O5- subset 2 isolates.

|  | Retail Meat | Clinical |  |  |  |  | Retail Meat |  |  | Clinical |
| --- | --- | --- | --- | --- | --- | --- | --- | --- | --- | --- |
|  | SC-29 | SC-30 | SC-31 | SC-32 | SC-33 | SC-34 | SC-35 | SC-36 | SC-37 | SC-38 |
| SC-29 | 0 |  |  |  |  |  |  |  |  |  |
| SC-30 | 58 | 0 |  |  |  |  |  |  |  |  |
| SC-31 | 61 | 57 | 0 |  |  |  |  |  |  |  |
| SC-32 | 27 | 57 | 60 | 0 |  |  |  |  |  |  |
| SC-33 | 72 | 66 | 73 | 73 | 0 |  |  |  |  |  |
| SC-34 | 71 | 67 | 70 | 70 | 83 | 0 |  |  |  |  |
| SC-35 | 28 | 58 | 61 | 21 | 74 | 71 | 0 |  |  |  |
| SC-36 | 31 | 61 | 64 | 24 | 77 | 74 | 17 | 0 |  |  |
| SC-37 | 35 | 65 | 68 | 28 | 81 | 78 | 21 | 24 | 0 |  |
| SC-38 | 24 | 52 | 55 | 23 | 68 | 65 | 24 | 27 | 31 | 0 |

### Genetic AMR profile generally aligned with observed AMR phenotype in each subset

To further characterize these isolates, assess whether they carry the same AMR genes, and determine AMR phenotype/genotype correlations, BARRGD and ResFinder 3.0 were used. Within the *S*. Heidelberg subset, six AMR genes were identified: *aadA1* (STR^r^), *aac(3)-IId* (GEN^r^), *aph(3’’)-Ib/strA* (STR^r^), *bla*_TEM-1B_ (broad-spectrum β-lactam^r^), *tet(A)* (TET^r^), and*fosA7* (fosfomycin; FOF^r^) (Fig. 4A). All six of these genes, except for *aph(3’’)-Ib/strA*, were identified in the assembled genomes of all isolates; only isolates SH-01 through SH-03, SH-06, SH-07, SH-09, and SH-10 carried *aph(3’’)-Ib/strA* (Fig. 4A). Interestingly, the genome of SH-04, when sequenced by PacBio, housed an additional copy of *bla*_TEM-1B_ (Fig. 4A). Indeed, *bla*_TEM-1B_ was located in a relatively short contig in the draft genomes of all isolates, likely indicating it did not assemble well, providing a possible explanation for why it was detected once in the MiSeq-based assembly, but twice in the PacBio-based genome. No known AMR-conferring chromosomal mutations were identified in any isolate within this subset or in either *S*. Typhimurium var. O5- subset.

**Figure 4.**
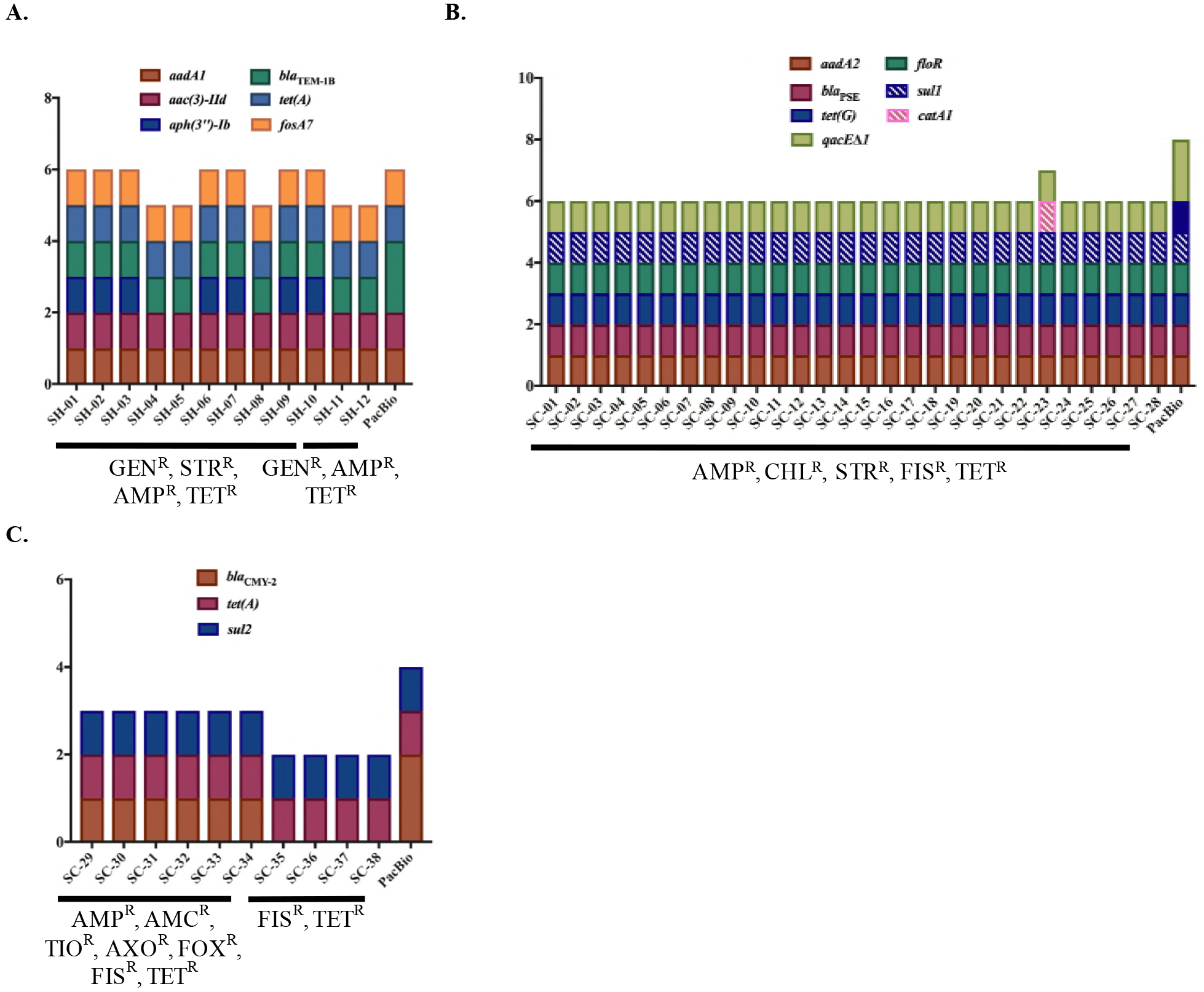
Identification of genetic resistance determinants in each subset. (A) *S*. Heidelberg subset. (B) *S*. Typhimurium var. O5- subset 1. (C) *S*. Typhimurium var. O5- subset 2. Each column represents the resistance genes identified in each isolate’s assembled genome, with the exception of the last column in each graph, which represents the genes identified in the chromosome and plasmid(s) of each isolate sequenced by PacBio: SH-04 in (A), SC-09 in (B), and SC-31 in (C). A combination of BARRGD (Accession number PRJNA313047; accessed April 2018) and ResFinder 3.0 (44) databases were used to identify genes. Solid-colored genes were present at > 99.3% nucleotide identity and 100% query coverage. Pattern-colored genes were present at > 99.3% nucleotide identity and > 63% query coverage. Each isolate’s phenotypic drug resistance profile is noted along the bottom. AMP=ampicillin; AMC=amoxicillin-clavulanic acid; TIO=ceftiofur; AXO=ceftriaxone; FOX=cefoxitin; CHL=chloramphenicol; GEN=gentamicin; STR=streptomycin; FIS=sulfisoxazole; TET=tetracycline.

Within *S*. Typhimurium var. O5- subset 1, a total of six different AMR genes were identified: *aadA2* (STR^r^), *bla*_pSE_/*bla*_CARB-2_ (broad-spectrum β-lactam^r^), *tet(G)* (TET^r^), *floR* (CHL^r^), *sull* (FIS^r^), and *catAl* (CHL^r^) (Fig. 4B). In addition, *qacEΔ*1, a gene that confers resistance to quaternary ammonium compounds was identified in all isolates (Fig. 4B). All AMR genes, but *catA1*, were identified in each isolate; *catA1* was only present in clinical isolate SC-23 (Fig. 4B). Within the PacBio genome of SC-09, an additional copy of *qacEAl* was identified, as well as a partial and complete copy of *sull*, which was only partially present once (~66% query coverage) in the assembled genome of each isolate (Fig. 4B).

Within the last subset, *S*. Typhimurium var. O5- subset 2, three AMR genes were identified: *bla*_CMY-2_ (broad and extended-spectrum β-lactam^r^), *tet(A)* (TET^r^), and *sul2* (FIS^r^), (Fig. 4C). Each AMR gene, except *bla*_CMY-2_, was identified in each assembled genome; *bla*_CMY-2_ was only identified in isolates SC-29 through SC-34 (Fig. 4C). Moreover, within the PacBio genome of SC-31, two *bla*_CMY-2_ genes were identified (Fig. 4C).

Accordingly, across all three subsets, there was a 100% correlation between AMR phenotype and genotype for β-lactam, gentamicin (an aminoglycoside), chloramphenicol, sulfonamide, and tetracycline antimicrobials; whereas, there was a 95% correlation for streptomycin (an aminoglycoside) (Table 5). Additionally, no correlation for fosfomycin could be calculated, as resistance to this drug was not phenotypically tested for.

**Table 5.**
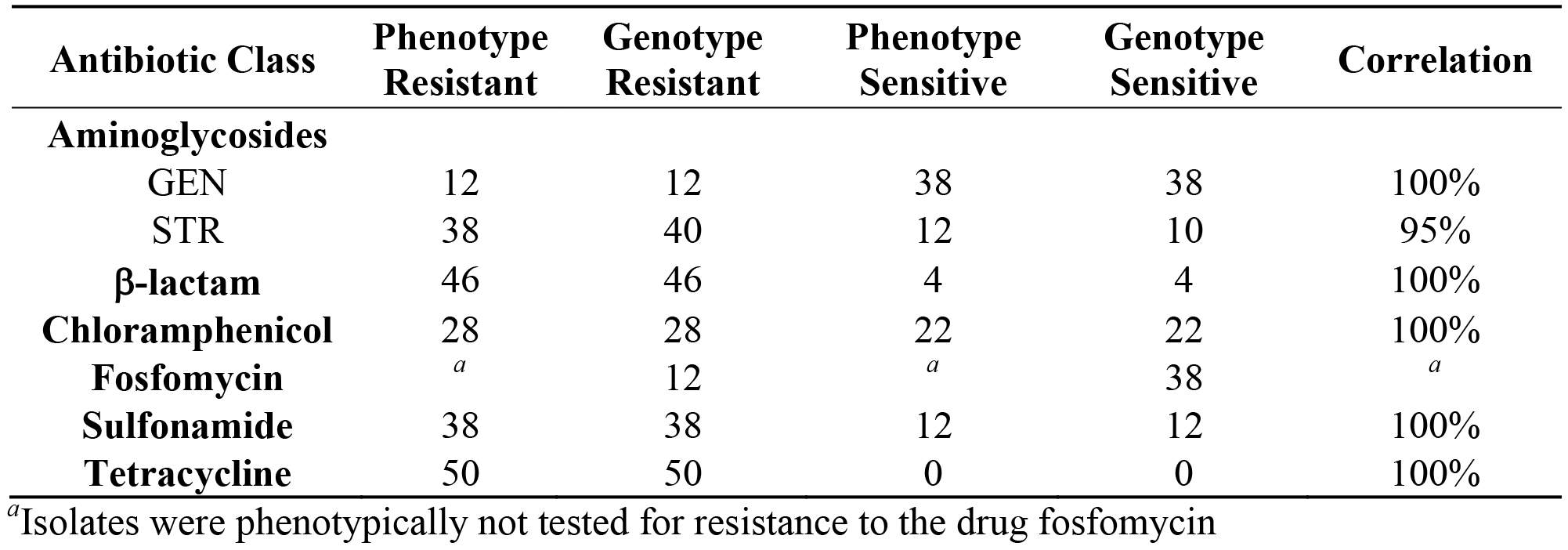
Overall correlation between AMR phenotype and genotype across all subsets.

### Identification of plasmid and chromosomal AMR determinants in each subset

Lastly, to further elucidate the genetic AMR profile of each isolate, plasmids were identified in an effort to determine and compare AMR gene location in each isolate. PacBio sequencing of SC-09, from *S*. Typhimurium var. O5- subset 1, revealed one plasmid with the replicon sequences, IncFIB(S), IncFII(S). This plasmid (pSC-09-1) was conserved in all isolates within this subset and was found to house multiple virulence-associated genes, but no AMR genes were identified (Fig. S3). This observation indicated that AMR was chromosomally carried in this subset. Indeed, all AMR genes identified in this subset were located on the SC-09 chromosome. Furthermore, following alignment, it was found that all isolates carried their AMR genes within an approximate 12-kb chromosomal region (Fig. 5). Moreover, annotation demonstrated that approximately 3-kb upstream of *sull* is an integrase and immediately downstream of *aadA2* is an additional integrase, suggesting that all AMR genes are associated with a similar mobile genetic element in this subset.

**Figure 5.**
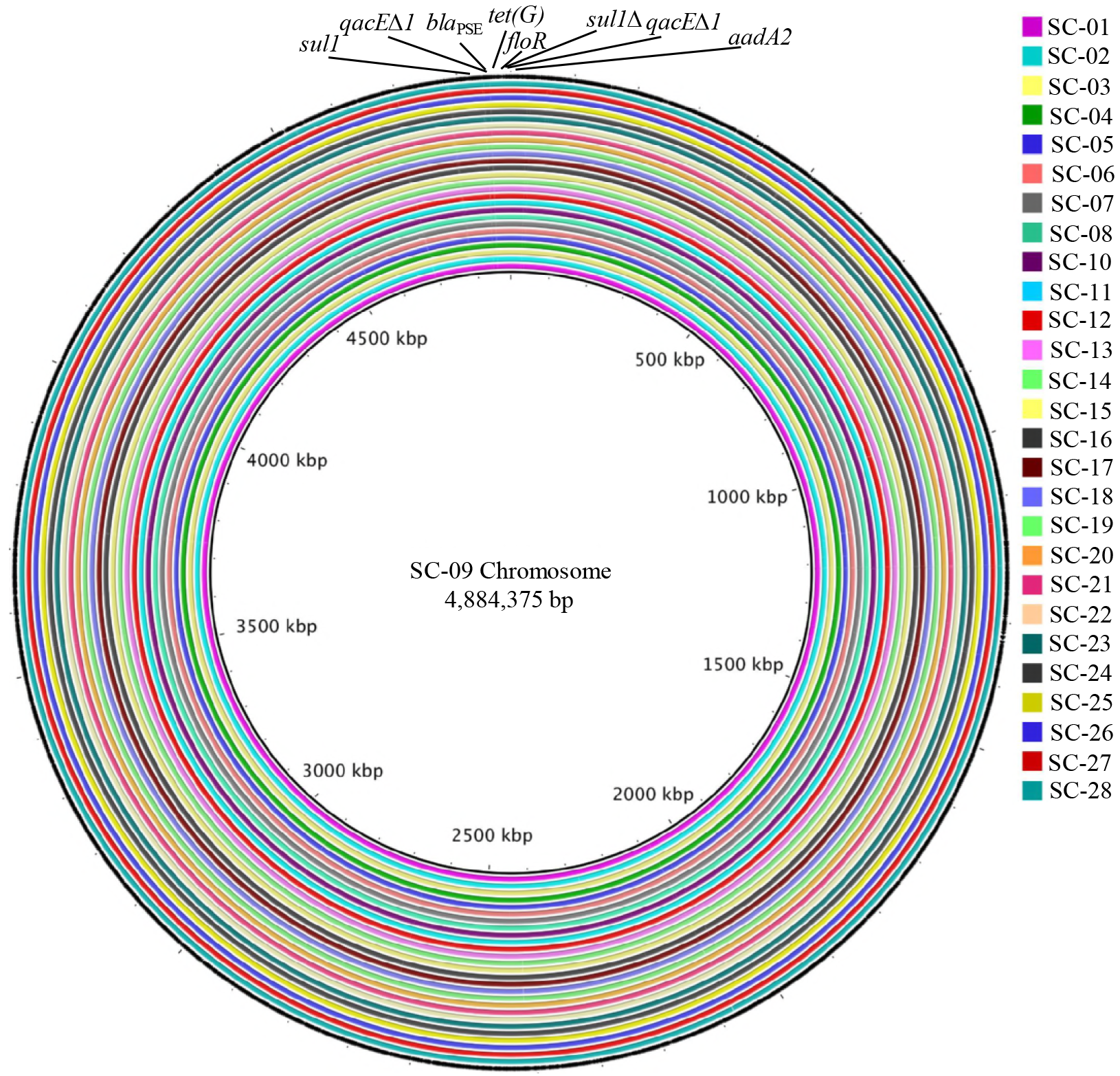
Chromosomally carried AMR in *S*. Typhimurium var. O5- subset 1. Alignment of the SC-09 chromosome, closed by PacBio sequencing, against the assembled genomes of SC-01 through SC-08 and SC-10 through SC-28 using BRIG (46). Each colored concentric ring represents one assembled genome aligning to the reference genome. The outermost ring represents open reading frames (ORFs) within the SC-09 chromosome and AMR gene locations are indicated.

Within the *S*. Heidelberg subset, PacBio sequencing of SH-04 resulted in two plasmids, pSH-04-1 (IncI1-alpha) and pSH-04-2 (IncX1). pSH-04-1 was present in both retail meat and all ten clinical isolates and housed four (*aac(3)-IId*, *bla*_TEM-1B_, *aadA1*, and *tet(A)*) of the six identified AMR genes (Fig. 6A). pSH-04-2 was also conserved in all twelve isolates and carried an additional copy of *bla*_TEM-1B_ (Fig. 6B). Interestingly, a BLAST search indicated that pSH-04-2 was different than other known *Salmonella* plasmids, with the top three hits sharing 98-99% nucleotide identity and 60-95% query coverage (pFDAARGOS_312_2, pSE95-0621-1, and pSTY3-1898) (Fig. S4A; Table S2); the main differences between the plasmids in Fig. S4A were the presence/absence of various mobile element and hypothetical proteins. Additionally, through annotation of the SH-04 chromosome, it was determined that *fosA7* was chromosomally-carried.

**Figure 6.**
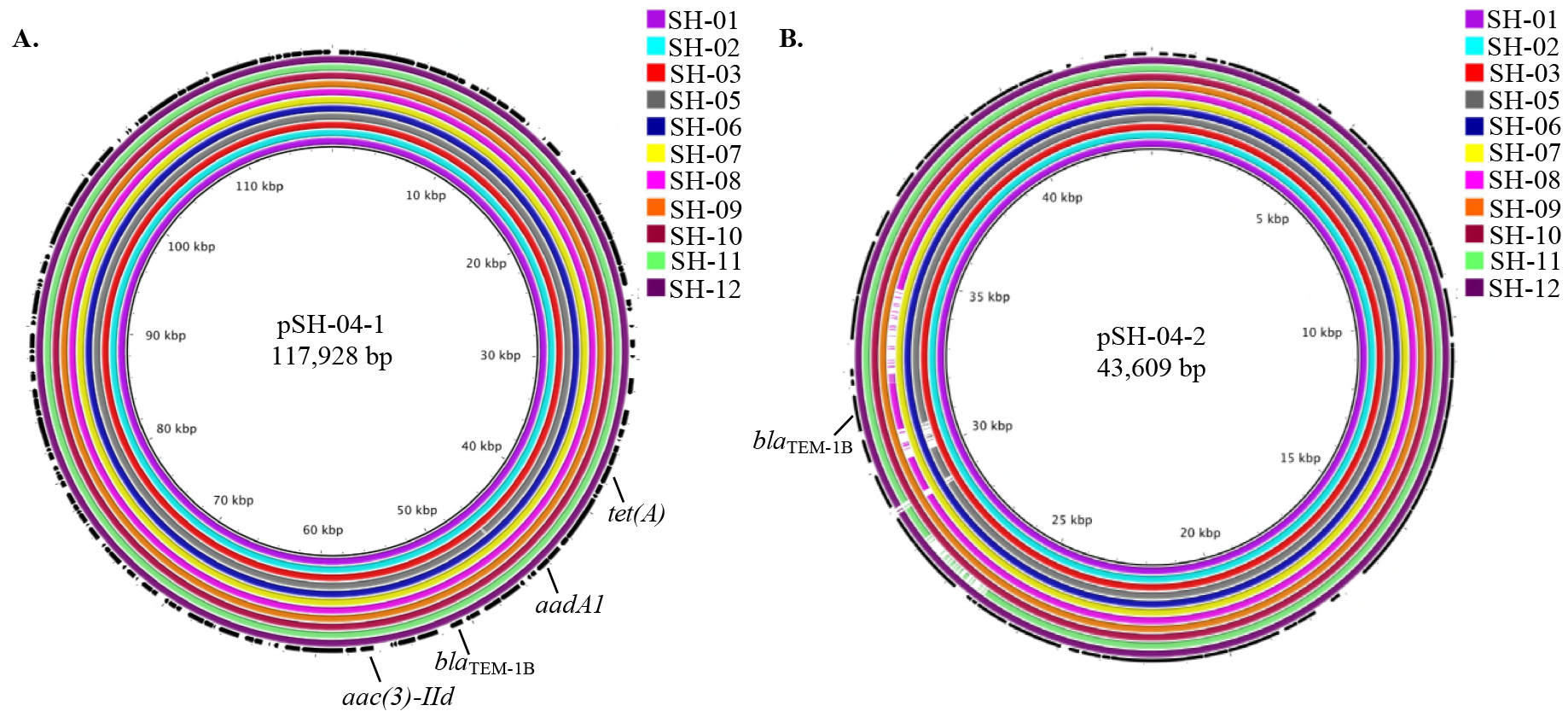
Comparative plasmid analysis in the *S*. Heidelberg subset. (A) Alignment of pSH-04-1 (IncI1-alpha), closed by PacBio sequencing, against the assembled genomes of SH-01 through SH-03 and SH-05 through SH-12 using BRIG (46). (B) Alignment of pSH-04-2 (IncX1), closed by PacBio sequencing, against the assembled genomes of SH-01 through SH-03 and SH-05 through SH-12. In (A) and (B), each colored concentric ring represents one assembled genome aligning to pSH-04-1 or pSH-04-2. The outermost ring in each panel represents ORFs in pSH-04-1 or pSH-04-2; the AMR annotations are included.

Finally, sequencing of SC-31, from *S*. Typhimurium var. O5- subset 2, resulted in three complete plasmid sequences. pSC-31-1 (IncI1-alpha) housed the beta-lactamase, *bla*_CMY-2_. Moreover, comparative sequence analysis demonstrated that pSC-31-1 was only present in isolates SC-29 through SC-33 (Fig. 7A). This result aligned clearly with phenotype as SC-29 through SC-33 were resistant to β-lactams, but SC-35 through SC-38 were sensitive. However, SC-34 also displayed resistance to β-lactam antibiotics, but did not appear to have this plasmid, despite carrying the *bla*_CMY-2_ gene (Fig. 4C). Therefore, it is postulated that SC-34 carries this gene on its chromosome. Supporting this hypothesis, SC-31 also carried a chromosomal copy of *bla*_CMY-2_. Conversely, pSC-31-2 (IncA/C2) was present in all isolates within this subset (Fig. 7B) and was found to house the remaining two AMR genes, *sul2* and *tet(A)*. Furthermore, a BLAST search revealed that only two other plasmids have been sequenced that show high similarity to this plasmid: pCFSAN001921 (99% nucleotide identity/100% query coverage) and pFDAARGOS_312_3 (99% nucleotide identity/92% query coverage) (Fig. S4B). Interestingly, both of those plasmids were also isolated from *S*. Typhimurium var. O5-. Lastly, pSC-31-3 (ColpVC) was only fully present in SC-29, SC-31, and SC-34, and housed no known AMR determinants (Fig. 7C).

**Figure 7.**
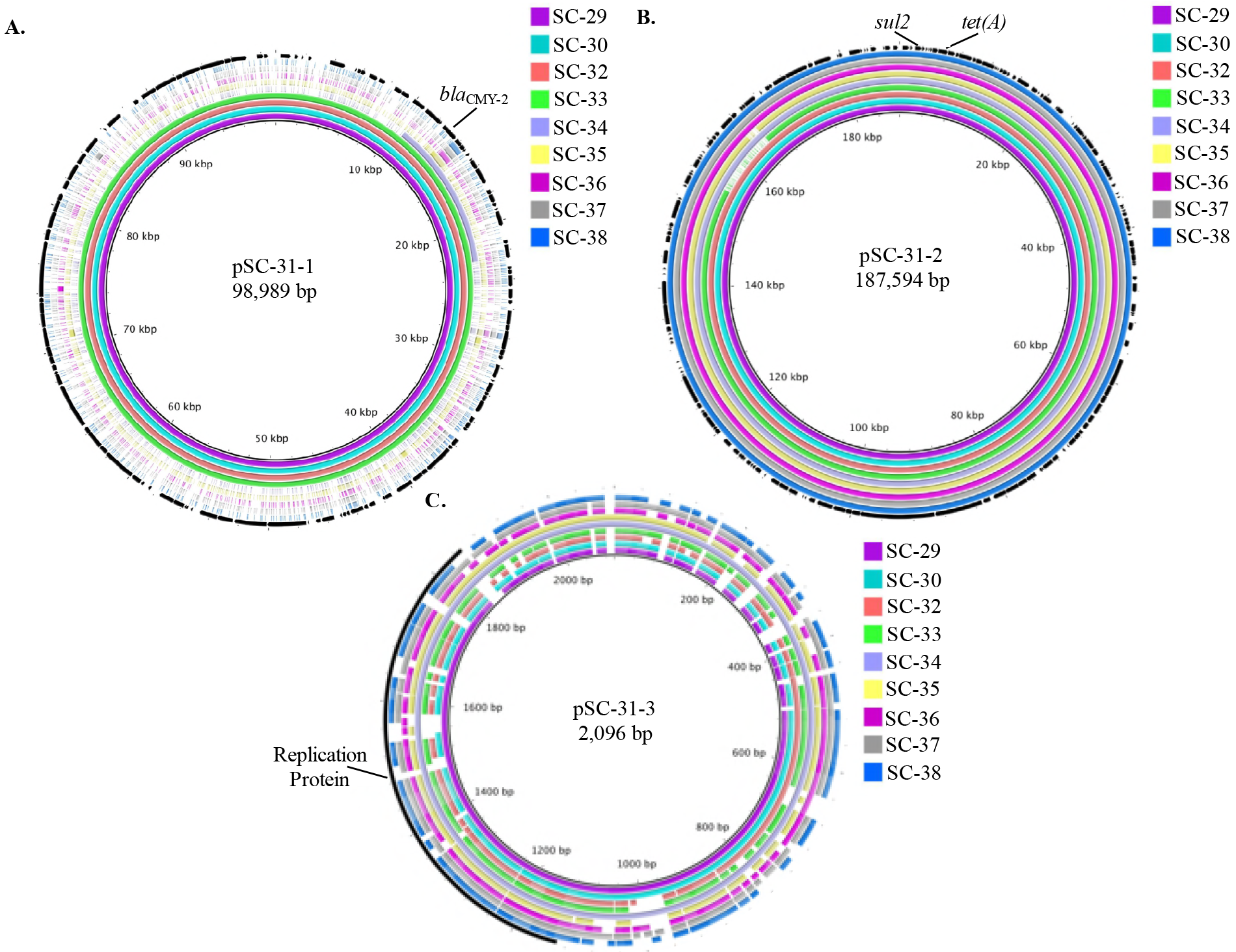
Comparative plasmid analysis in *S*. Typhimurium var. 05- subset 2. (A) Alignment of pSC-31-1 (Incll-alpha), closed by PacBio sequencing, against the assembled genomes of SC-29, SC-30, and SC-32 through SC-38 using BRIG (46). The assembled genomes of SC-34 through SC-38 did not align to the reference plasmid. (B) Alignment of pSC-31-2 (IncA/C2), closed by PacBio sequencing, against the assembled genomes of SC-29, SC-30, and SC-32 through SC-38. (C) Alignment of pSC-31-3 (ColpVC), closed by Illumina sequencing, against the assembled genomes of SC-29, SC-30, and SC-32 through SC-38. In (A-C), each colored concentric ring represents one assembled genome aligning to either pSC-31-1, pSC-31-2, or pSC-31-3. The outermost ring in each panel represents ORFs in each plasmid; the AMR annotations are included.

## Discussion

An important source of *Salmonella* is retail meat; in Pennsylvania, previous studies have demonstrated that non-typhoidal *Salmonella*, including isolates that are drug-resistant, are prevalent on meat products sold at grocery stores and farmers’ markets (48, 49). Here, we exploited the strength of WGS to compare the genetic characteristics of three historic subsets of drug-resistant and PFGE-matched retail meat and clinical *S*. Heidelberg and *S*. Typhimurium var. O5- isolates, to reassess their relatedness and identify their resistome.

Two previous studies have used WGS to compare S. Heidelberg isolated from multiple sources, including humans and retail meats. Hoffmann *et al*. (14) utilized 454 sequencing and SNP analysis to distinguish outbreak-associated Heidelberg isolates from non-outbreak isolates, collected from several sources between 1982 and 2011, with similar PFGE patterns. Edirmanasinghe *et al*. (24) used WGS to characterize *S*. Heidelberg from various sources, collected through routine surveillance in Canada. However, in this study, we focused on directly comparing the genomic characteristics of 50 historic retail meat and human S. Heidelberg and S. Typhimurium var. O5- isolates that matched, importantly, by multiple conventional methods (PFGE and AST) and were collected through routine surveillance in Pennsylvania between 2009 and 2014. The narrow geographical region and temporal distribution of this collection is reflective of what other state public health laboratories would examine during their own retrospective and comparative analyses using WGS.

### Comparative SNP analysis in *S*. Heidelberg subset reveals isolates are closely related

Within the *S.* Heidelberg subset, comparative SNP analysis revealed that the retail meat isolates, SH-01 and SH-02, were separated by 6 to 12 SNPs from the ten matched clinical isolates (Table 2). Prior studies have reported SNP differences of 0 to 4, an average of 17, and 4 to 19 for previous confirmed S. Heidelberg outbreaks (11, 14, 19). Thus, the observed SNP distances between the retail meat and clinical isolates within this subset align with outbreak-associated values reported in the literature for the Heidelberg serovar, indicating that these isolates are closely related and may share a common source. However, due to the historic nature of these isolates, an epidemiological link was not investigated between the food and human isolates in this case. Moreover, the retail meat isolates were collected in 2013, whereas, the majority of the clinical isolates were isolated from 2010 and 2011. Accordingly, these data underscore the importance of interpreting genomic results in the context of epidemiological data, which is particularly crucial when analyzing historic isolate collections.

### Comparative SNP analysis in S. Typhimurium var. O5- subsets reveals genetic differences

Conversely, comparative SNP analysis revealed a range of phylogenetic relationships within each S. Typhimurium var. O5- subset (Tables 3, 4). Importantly, the retail meat isolate within subset 1 was separated from all clinical isolates by 41 to 96 SNPs (Table 3); similarly, the four retail meat isolates within subset 2 were separated from the six clinical isolates by 21 to 81 SNPs (Table 4). Previous reports have determined that S. Typhimurium outbreak-associated isolates have been separated by 2 to 12, 3 to 30, 0 to 12, 0 to 7, and a maximum of 3 SNPs (19, 22, 50, 51). Furthermore, others have proposed specific SNP cutoffs to classify an isolate as outbreak-associated (22); however, a clearly defined consensus in the field on a maximum SNP distance threshold for outbreak analysis has not been established. Nonetheless, our data suggest that there is not a genetic link between the retail meat and clinical isolates within the two *S*. Typhimurium var. O5- subsets, despite them matching by conventional methods. Indeed, previous studies have found similar genetic distances between PFGE-matched sporadic and outbreak isolates; for example, one sporadic *S*. Bareilly isolate that shared the same primary PFGE pattern as an S. Bareilly outbreak was separated by 117 SNPs from those strains (intra-outbreak SNP distance was 1 to 6) (20).

### AMR phenotype and resistome were well correlated in each subset

Within this study, we also determined the resistome of each isolate, which included the identification of specific AMR genes and elucidation of each gene’s genomic context, in an effort to assess the role of WGS in enhancing integrated surveillance for drug-resistant *Salmonella* in Pennsylvania. Overall, the AMR profiles, as determined by conventional AST, correlated well with AMR genotype. Across all subsets, a 100% correlation was observed for all tested antimicrobials, except streptomycin, where a 95% correlation was observed (Table 5). Similarly, in a comprehensive *Salmonella* study encompassing 640 isolates, the AMR phenotype/genotype correlation was 99% (13); these results are corroborated by a smaller study that observed a 100% correlation between AMR phenotype and genotype for 49 *Salmonella* isolates from swine (52). Furthermore, the slightly lower streptomycin correlation that we observed has been noted previously. McDermott *et al*. (13) postulate that this discordance between phenotype and genotype is likely the result of a MIC breakpoint value that is too high. Consequently, another study found that by lowering the breakpoint MIC value for streptomycin resistance from ≥ 64 μg/L to ≥ 32 μg/L, a higher correlation was observed (53). Thus, it is plausible that the lower correlation observed in this study is also the result of not enough isolates being considered resistant based on standard MIC testing; however, silent genes or a non-functional protein product could also be responsible.

### AMR is primarily plasmid-mediated in the *S*. Heidelberg subset

Within the S. Heidelberg subset, six AMR genes were identified (Fig. 4A). These genes or their close variants have been identified in S. Heidelberg previously (14, 54–58). Specifically, *fosA7* was identified in *Salmonella* for the first time in 2017; this particular gene sequence was only found in 35 of the approximately 40,000 *Salmonella* draft and complete genomes in NCBI, of which 75% were S. Heidelberg (58). Notably, *fosA7* was located in both retail meat and all ten clinical S. Heidelberg isolates in this study. Moreover, PacBio sequencing revealed that this gene was located on the chromosome of SH-04, consistent with prior data suggesting that *fosA7* is exclusively chromosomal in *Salmonella* (58). PacBio sequencing also elucidated the presence of multi-copy AMR genes, which were not detected in the draft genomes within this subset (i.e. *bla*_TEM-1B_) and in each S. Typhimurium var. O5- subset as well; this observation highlights a potential advantage of incorporating long-read sequencing into routine surveillance methods.

In addition, comparative plasmid analysis identified two plasmids that were present in each retail meat and clinical isolate. pSH-04-1, an IncI1-alpha plasmid, carried four of the six AMR genes (*aadA1, aac(3)-IId*, *bla*_TEM-1B_, *tet(A)*) (Fig. 6A). Similar resistance genes have been reported on an IncI1 plasmid in S. Heidelberg previously (14). This replicon type plasmid has also been found to house sulfonamide resistance and importantly, the **bla*_CMY_* beta-lactamase, which encodes resistance to extended-spectrum cephalosporins, in *S*. Heidelberg (24, 57, 59). pSH-04-2, an IncX1 plasmid, housed an additional copy of *bla*_TEM-1B_ (Fig. 6B); IncX1 plasmids have been identified in *S*. Heidelberg isolates previously (14).

### AMR is chromosomally-carried in *S*. Typhimurium var. O5- subset 1

Within S. Typhimurium var. O5- subset 1, seven different resistance genes were identified (Fig. 4B). Annotation and subsequent alignment of each isolate’s chromosome determined that all of the identified genes, with the exception of *catA1* in isolate SC-23, were located within a 12-kb chromosomal region (Fig. 5). This AMR gene topology is typical of *Salmonella* Typhimurium strains that also display the ACSSuT penta-resistance pattern. Previous work has demonstrated that a region of the chromosome, termed *Salmonella* genomic island 1 (SGI1), houses the AMR gene cluster that is responsible for this phenotype; within this region, the AMR genes, *floR* and *tet(G)*, are flanked by integrons carrying the AMR genes, *aadA2* and *bla*_PSE_/*bla*_CARB-2_ (60, 61). Consistent with this observation, the single plasmid identified within this subset did not carry AMR genes.

### AMR is plasmid-mediated in S. Typhimurium var. O5- subset 2

Lastly, within S. Typhimurium var. O5- subset 2, three AMR genes were identified (Fig. 4C). These genes have previously been identified in *S*. Typhimurium and *S*. Typhimurium var. O5- (62–64). Comparative plasmid analysis revealed that pSC-31-1 (IncI1-alpha), which carried *bla*_CMY_-_2_, was present only in isolates SC-29 through SC-33 (Fig. 7A). Indeed, IncI1 plasmids are frequently associated with this gene in *Salmonella*, including *S*. Typhimurium var. O5- (62). Moreover, this genetic observation is consistent with the AST results, as isolates SC-29 through SC-33 were resistant to β-lactams, whereas, SC-35 through SC-38 were sensitive. Intriguingly, isolate SC-34 was resistant to β-lactams and was found to carry the *bla*_CMY-2_ gene; however, the IncI1-alpha plasmid was not present in this isolate. Indeed, a previous S. Heidelberg study identified *bla*_CMY-2_ on the chromosome, which is suggestive of plasmid integration being possible (24). Conversely, the second plasmid, pSC-31-2, was found in all retail meat and clinical isolates within this subset (Fig. 7B). This 188 kb IncA/C2 plasmid housed the AMR genes, *sul2* and *tet(A)*. Notably, a BLAST query indicated that this plasmid was only similar to two other plasmids, both of which were from other S. Typhimurium var. O5- isolates (Fig. S4B). When comparing these plasmids to pSC-31-2, the main differences were the presence or absence of multiple conjugal transfer-associated and hypothetical proteins. Accordingly, these data suggest that pSC-31-2 is a novel version of an IncA/C2, AMR-encoding plasmid that appears to generally be restricted to the serovar Typhimurium var. O5-.

In summary, this study demonstrated that historic retail meat and human *Salmonella* isolates, collected through routine monitoring, that are indistinguishable by the conventional methods PFGE and AST, could be different strains, further underscoring the power of WGS. These data also highlight the importance and necessity of interpreting WGS data in the context of epidemiological findings—a point that is particularly crucial, when analyzing historic isolate collections. In addition, we evaluated the role of WGS in enhancing integrated surveillance of drug-resistant *Salmonella* from retail meat and clinical sources in Pennsylvania, by identifying resistance genes and characterizing their genomic environment, as this information is vital to understanding how resistance is disseminating. We observed that resistance phenotype and genotype correlated well in each isolate and that the same AMR-encoding plasmids were found in both the retail meat and clinical isolates. As one of the first studies to directly compare historic retail meat and clinical *Salmonella* isolates using WGS, these results demonstrate the usefulness and value of WGS to public health laboratories performing retrospective comparisons of bacterial isolates from multiple sources.

## Acknowledgements

We thank Lisa Dettinger, James Tait, Carina Davis, Melinda Johnston, and Barry Perry for assistance with preparation of *Salmonella* isolates. A.B.K. was supported by The Black Endowed Graduate Fellowship through the Penn State College of Agricultural Sciences. This work was supported by the U.S. Food and Drug Administration grant number 1U18FD006222-01 to E.G.D. for support of GenomeTrakr in Pennsylvania, the U.S. Food and Drug Administration grant number 1U01FD006253-01 to PADOH for collaboration in the National Antimicrobial Resistance Monitoring System, Centers for Disease Control and Prevention Epidemiology and Laboratory Capacity (ELC) for collaboration in National Antimicrobial Resistance Monitoring (CDC-RFA-CI10-101204PPHF13), and the USDA National Institute of Food and Agriculture Federal Appropriations under Project PEN04522 and Accession number 0233376.

## References

1. Scallan E, Hoekstra RM, Angulo FJ, Tauxe RV, Widdowson M-A, Roy SL, Jones JL, Griffin PM. 2011. Foodborne illness acquired in the United States—major pathogens. Emerg Infect Dis 17:7–15.

2. CDC. 2013. Antibiotic resistance threats in the United States, 2013. Atlanta, Georgia. https://www.cdc.gov/drugresistance/threat-report-2013/pdf/ar-threats-2013-508.pdf

3. Jackson BR, Griffin PM, Cole D, Walsh KA, Chai SJ. 2013. Outbreak-associated Salmonella enterica serotypes and food commodities, United States, 1998-2008. Emerg Infect Dis 19:1239–1244.

4. CDC. 2017. Surveillance for foodborne disease outbreaks, United States, 2015, annual report. Atlanta, Georgia. https://www.cdc.gov/foodsafety/pdfs/2015FoodBorne0utbreaks_508.pdf

5. Boore AL, Hoekstra RM, Iwamoto M, Fields PI, Bishop RD, Swerdlow DL. 2015. Salmonella enterica infections in the United States and assessment of coefficients of variation: a novel approach to identify epidemiologic characteristics of individual serotypes, 1996-2011. PLoS One 10:e0145416.

6. FDA. 2015. 2014-2015 Retail meat interim report. Rockville, MD. https://www.fda.gov/downloads/AnimalVeterinary/SafetyHealth/AntimicrobialResistance/NationalAntimicrobialResistanceMonitoringSystem/UCM498134.pdf

7. Acheson D, Hohmann EL. 2001. Nontyphoidal salmonellosis. Clin Infect Dis 32:263–269.

8. White House. 2014. National Strategy For Combating Antibiotic-Resistant Bacteria. Washington DC. https://obamawhitehouse.archives.gov/sites/default/files/docs/carb_national_strategy.pdf

9. Gilbert JM, White DG, McDermott PF. 2007. The US National Antimicrobial Resistance Monitoring System. Future Microbiol 2:493–500.

10. Salipante SJ, SenGupta DJ, Cummings LA, Land TA, Hoogestraat DR, Cookson BT. 2015. Application of whole-genome sequencing for bacterial strain typing in molecular epidemiology. J Clin Microbiol 53:1072–1079.

11. Bekal S, Berry C, Reimer AR, Van Domselaar G, Beaudry G, Fournier E, Doualla-Bell F, Levac E, Gaulin C, Ramsay D, Huot C, Walker M, Sieffert C, Tremblay C. 2016. Usefulness of high-quality core genome single-nucleotide variant analysis for subtyping the highly clonal and the most prevalent Salmonella enterica serovar Heidelberg clone in the context of outbreak investigations. J Clin Microbiol 54:289–295.

12. Kahlmeter G. 2014. Defining antibiotic resistance-towards international harmonization. Ups J Med Sci 119:78–86.

13. McDermott PF, Tyson GH, Kabera C, Chen Y, Li C, Folster JP, Ayers SL, Lam C, Tate HP, Zhao S. 2016. Whole-genome sequencing for detecting antimicrobial resistance in Nontyphoidal Salmonella. Antimicrob Agents Chemother 60:5515–5520.

14. Hoffmann M, Zhao S, Pettengill J, Luo Y, Monday SR, Abbott J, Ayers SL, Cinar HN, Muruvanda T, Li C, Allard MW, Whichard J, Meng J, Brown EW, McDermott PF. 2014. Comparative genomic analysis and virulence differences in closely related Salmonella enterica serotype Heidelberg isolates from humans, retail meats, and animals. Genome Biol Evol 6:1046–1068.

15. Phillips A, Sotomayor C, Wang Q, Holmes N, Furlong C, Ward K, Howard P, Octavia S, Lan R, Sintchenko V. 2016. Whole genome sequencing of Salmonella Typhimurium illuminates distinct outbreaks caused by an endemic multi-locus variable number tandem repeat analysis type in Australia, 2014. BMC Microbiol 16.

16. den Bakker HC, Allard MW, Bopp D, Brown EW, Fontana J, Iqbal Z, Kinney A, Limberger R, Musser KA, Shudt M, Strain E, Wiedmann M, Wolfgang WJ. 2014. Rapid whole-genome sequencing for surveillance of Salmonella enterica serovar Enteritidis. Emerg Infect Dis 20:1306–1314.

17. Allard MW, Luo Y, Strain E, Li C, Keys CE, Son I, Stones R, Musser SM, Brown EW. 2012. High resolution clustering of Salmonella enterica serovar Montevideo strains using a next-generation sequencing approach. BMC Genomics 13:32.

18. Wilson MR, Brown E, Keys C, Strain E, Luo Y, Muruvanda T, Grim C, Jean-Gilles Beaubrun J, Jarvis K, Ewing L, Gopinath G, Hanes D, Allard MW, Musser S. 2016. Whole genome DNA sequence analysis of Salmonella subspecies enterica serotype Tennessee obtained from related peanut butter foodborne outbreaks. PLoS One 11:e0146929.

19. Leekitcharoenphon P, Nielsen EM, Kaas RS, Lund O, Aarestrup FM. 2014. Evaluation of whole genome sequencing for outbreak detection of Salmonella enterica. PLoS One 9:e87991.

20. Hoffmann M, Luo Y, Monday SR, Gonzalez-Escalona N, Ottesen AR, Muruvanda T, Wang C, Kastanis G, Keys C, Janies D, Senturk IF, Catalyurek U V., Wang H, Hammack TS, Wolfgang WJ, Schoonmaker-Bopp D, Chu A, Myers R, Haendiges J, Evans PS, Meng J, Strain EA, Allard MW, Brown EW. 2016. Tracing origins of the Salmonella Bareilly strain causing a food-borne outbreak in the United States. J Infect Dis 213:502–508.

21. Taylor AJ, Lappi V, Wolfgang WJ, Lapierre P, Palumbo MJ, Medus C, Boxrud D. 2015. Characterization of foodborne outbreaks of Salmonella enterica serovar Enteritidis with whole-genome sequencing single nucleotide polymorphism-based analysis for surveillance and outbreak detection. J Clin Microbiol 53:3334–3340.

22. Octavia S, Wang Q, Tanaka MM, Kaur S, Sintchenko V, Lan R. 2015. Delineating community outbreaks of Salmonella enterica serovar Typhimurium by use of whole-genome sequencing: insights into genomic variability within an outbreak. J Clin Microbiol 53:1063–1071.

23. CDC. 2014. NARMS 2014 human isolates surveillance report. Atlanta, Georgia. https://www.cdc.gov/narms/pdf/2014-annual-report-narms-508c.pdf

24. Edirmanasinghe R, Finley R, Parmley EJ, Avery BP, Carson C, Bekal S, Golding G, Mulvey MR. 2017. A whole genome sequencing approach to study cefoxitin-resistant Salmonella enterica serovar Heidelberg from various aources. Antimicrob Agents Chemother 61:e01919–e16.

25. Zhao S, Tyson GH, Chen Y, Li C, Mukherjee S, Young S, Lam C, Folster JP, Whichard JM, McDermott PF. 2015. Whole-genome sequencing analysis accurately predicts antimicrobial resistance phenotypes in Campylobacter spp. Appl Environ Microbiol 82:459–466.

26. FDA. 2012. 2012 Retail meat report National Antimicrobial Resistance Monitoring System. Laurel, Maryland. https://www.fda.gov/downloads/animalveterinary/safetyhealth/antimicrobialresistance/nationalantimicrobialresistancemonitoringsystem/ucm442212.pdf

27. Pennsylvania Department of Health. List of reportable diseases. http://www.health.pa.gov/Your-Department-of-Health/Offices%20and%20Bureaus/epidemiology/Pages/Reportable-Diseases.aspx#.Wy6PXBJKho6

28. CDC. 2017. Standard Operating Procedure for PulseNet PFGE of Escherichia coli 0157:H7, Escherichia coli non-0157 (STEC), Salmonella serotypes, Shigella sonnei and Shigella flexneri. Atlanta, GA. https://www.cdc.gov/pulsenet/pdf/ecoli-shigella-salmonella-pfge-protocol-508c.pdf

29. Sandt CH, Fedorka-Cray PJ, Tewari D, Ostroff S, Joyce K, Mikanatha NM. 2013. A comparison of non-typhoidal Salmonella from humans and food animals using pulsed-field gel electrophoresis and antimicrobial susceptibility patterns. PLoS One 8:e77836.

30. FDA. Breakpoints Table 1. interpretive criteria used for susceptibility testing of Salmonella and E. coli. https://www.fda.gov/downloads/AnimalVeterinary/SafetyHealth/AntimicrobialResistance/NationalAntimicrobialResistanceMonitoringSystem7UCM581395.pdf

31. Yao K, Muruvanda T, Roberts RJ, Payne J, Allard MW, Hoffmann M. 2016. Complete genome and methylome sequences of Salmonella enterica subsp. enterica serovar Panama (ATCC 7378) and Salmonella enterica subsp. enterica serovar Sloterdijk (ATCC 15791). Genome Announc 4:e00133–16.

32. Aziz RK, Bartels D, Best AA, DeJongh M, Disz T, Edwards RA, Formsma K, Gerdes S, Glass EM, Kubal M, Meyer F, Olsen GJ, Olson R, Osterman AL, Overbeek RA, McNeil LK, Paarmann D, Paczian T, Parrello B, Pusch GD, Reich C, Stevens R, Vassieva O, Vonstein V, Wilke A, Zagnitko O. 2008. The RAST Server: Rapid Annotations using Subsystems Technology. BMC Genomics 9:75.

33. Babraham Bioinformatics. 2016. FastQC v0.11.5.

34. Li H. 2013. Aligning sequence reads, clone sequences and assembly contigs with BWA-MEM. arXiv 0:1–3.

35. Li H, Handsaker B, Wysoker A, Fennell T, Ruan J, Homer N, Marth G, Abecasis G, Durbin R, 1000 Genome Project Data Processing Subgroup 1000 Genome Project Data Processing. 2009. The Sequence Alignment/Map format and SAMtools. Bioinformatics 25:2078–2079.

36. Bankevich A, Nurk S, Antipov D, Gurevich AA, Dvorkin M, Kulikov AS, Lesin VM, Nikolenko SI, Pham S, Prjibelski AD, Pyshkin A V, Sirotkin A V, Vyahhi N, Tesler G, Alekseyev MA, Pevzner PA. 2012. SPAdes: a new genome assembly algorithm and its applications to single-cell sequencing. J Comput Biol 19:455–477.

37. Gurevich A, Saveliev V, Vyahhi N, Tesler G. 2013. QUAST: quality assessment tool for genome assemblies. Bioinformatics 29:1072–1075.

38. Zhang S, Yin Y, Jones MB, Zhang Z, Deatherage Kaiser BL, Dinsmore BA, Fitzgerald C, Fields PI, Deng X. 2015. Salmonella serotype determination utilizing high-throughput genome sequencing data. J Clin Microbiol 53:1685–1692.

39. Petkau A, Mabon P, Sieffert C, Knox NC, Cabral J, Iskander M, Iskander M, Weedmark K, Zaheer R, Katz LS, Nadon C, Reimer A, Taboada E, Beiko RG, Hsiao W, Brinkman F, Graham M, Van Domselaar G. 2017. SNVPhyl: a single nucleotide variant phylogenomics pipeline for microbial genomic epidemiology. Microb Genomics 3.

40. Guindon S, Dufayard J-F, Lefort V, Anisimova M, Hordijk W, Gascuel O. 2010. New algorithms and methods to estimate maximum-likelihood phylogenies: assessing the performance of PhyML 3.0. Syst Biol 59:307–321.

41. Danecek P, Auton A, Abecasis G, Albers CA, Banks E, Depristo MA, Handsaker RE, Lunter G, Marth GT, Sherry ST, Mcvean G, Durbin R, Project G, Group A. 2011. The variant call format and VCFtools. Bioinforma Appl Note 27:2156–2158.

42. Cingolani P, Platts A, Wang LL, Coon M, Nguyen T, Wang L, Land SJ, Lu X, Ruden DM. 2012. A program for annotating and predicting the effects of single nucleotide polymorphisms, SnpEff: SNPs in the genome of Drosophila melanogaster strain w1118; iso-2; iso-3. Fly (Austin) 6:80–92.

43. Camacho C, Coulouris G, Avagyan V, Ma N, Papadopoulos J, Bealer K, Madden TL. 2009. BLAST+: architecture and applications. BMC Bioinformatics 10:421.

44. Zankari E, Hasman H, Cosentino S, Vestergaard M, Rasmussen S, Lund O, Aarestrup FM, Larsen MV. 2012. Identification of acquired antimicrobial resistance genes. J Antimicrob Chemother 67:2640–2644.

45. Carattoli A, Zankari E, Garcia-Fernandez A, Larsen MV, Lund O, Villa L, Aarestrup FM, Hasman H. 2014. PlasmidFinder and pMLST: in silico detection and typing of plasmids. Antimicrob Agents Chemother 58:3895–3903.

46. Alikhan N-F, Petty NK, Ben Zakour NL, Beatson SA. 2011. BLAST Ring Image Generator (BRIG): simple prokaryote genome comparisons. BMC Genomics 12:402.

47. Altschul SF, Gish W, Miller W, Myers EW, Lipman DJ. 1990. Basic local alignment search tool. J Mol Biol 215:403–410.

48. M’ikanatha NM, Sandt CH, Localio AR, Tewari D, Rankin SC, Whichard JM, Altekruse SF, Lautenbach E, Folster JP, Russo A, Chiller TM, Reynolds SM, Mcdermott PF. 2010. Multidrug-resistant Salmonella isolates from retail chicken meat compared with human clinical isolates. Foodborne Pathog Dis 7:929–934.

49. Scheinberg J, Doores S, Cutter CN. 2013. A microbiological comparison of poultry products obtained from farmers’ markets and supermarkets in Pennsylvania. J Food Saf 33:259–264.

50. Gymoese P, Sørensen G, Litrup E, Olsen JE, Nielsen EM, Torpdahl M. 2017. Investigation of outbreaks of Salmonella enterica serovar Typhimurium and its monophasic variants using whole-genome sequencing, Denmark. Emerg Infect Dis 23:1631–1639.

51. Ashton PM, Peters T, Ameh L, McAleer R, Petrie S, Nair S, Muscat I, de Pinna E, Dallmand T. 2015. Whole genome sequencing for the retrospective investigation of an outbreak of Salmonella Typhimurium DT 8. PLoS Curr 7.

52. Zankari E, Hasman H, Kaas RS, Seyfarth AM, Agerso Y, Lund O, Larsen M V., Aarestrup FM. 2013. Genotyping using whole-genome sequencing is a realistic alternative to surveillance based on phenotypic antimicrobial susceptibility testing. J Antimicrob Chemother 68:771–777.

53. Tyson GH, Li C, Ayers S, McDermott PF, Zhao S. 2016. Using whole-genome sequencing to determine appropriate streptomycin epidemiological cutoffs for Salmonella and Escherichia coli. FEMS Microbiol Lett 363.

54. Chen S, Zhao S, White DG, Schroeder CM, Lu R, Yang H, McDermott PF, Ayers S, Meng J. 2004. Characterization of multiple-antimicrobial-resistant Salmonella serovars isolated from retail meats. Appl Environ Microbiol 70:1–7.

55. Folster JP, Pecic G, Rickert R, Taylor J, Zhao S, Fedorka-Cray PJ, Whichard J, McDermott P. 2012. Characterization of multidrug-resistant Salmonella enterica serovar Heidelberg from a ground turkey-associated outbreak in the United States in 2011. Antimicrob Agents Chemother 56:3465–3466.

56. Patchanee P, Zewde BM, Tadesse DA, Hoet A, Gebreyes WA. 2008. Characterization of multidrug-resistant Salmonella enterica serovar Heidelberg isolated from humans and animals. Foodborne Pathog Dis 5:839–851.

57. Han J, Lynne AM, David DE, Tang H, Xu J, Nayak R, Kaldhone P, Logue CM, Foley SL. 2012. DNA sequence analysis of plasmids from multidrug resistant Salmonella enterica serotype Heidelberg isolates. PLoS One 7:e51160.

58. Rehman MA, Yin X, Persaud-Lachhman MG, Diarra MS. 2017. First detection of a fosfomycin resistance gene, fosA7, in Salmonella enterica serovar Heidelberg isolated from broiler chickens. Antimicrob Agents Chemother 61:e00410–e00417.

59. Folster JP, Pecic G, Singh A, Duval B, Rickert R, Ayers S, Abbott J, Mcglinchey B, Bauer-Turpin J, Haro J, Hise K, Zhao S, Fedorka-Cray PJ, Whichard J, Mcdermott PF. 2012. Characterization of extended-spectrum cephalosporin-resistant Salmonella enterica serovar Heidelberg isolated from food animals, retail meat, and humans in the United States 2009. Foodborne Pathog Dis 9:638–645.

60. Boyd DA, Peters GA, Ng L-K, Mulvey MR. 2000. Partial characterization of a genomic island associated with the multidrug resistance region of Salmonella enterica Typhymurium DT104. FEMS Microbiol Lett 189:285–291.

61. Boyd D, Peters GA, Cloeckaert A, Boumedine KS, Chaslus-Dancla E, Imberechts H, Mulvey MR. 2001. Complete nucleotide sequence of a 43-kilobase genomic island associated with the multidrug resistance region of Salmonella enterica serovar Typhimurium DT104 and its identification in phage type DT120 and serovar Agona. J Bacteriol 183:5725–5732.

62. Gray JT, Hungerford LL, Fedorka-Cray PJ, Headrick ML. 2004. Extended-spectrum-cephalosporin resistance in Salmonella enterica isolates of animal origin. Antimicrob Agents Chemother 48:3179–3181.

63. Frech G, Kehrenberg C, Schwarz S. 2003. Resistance phenotypes and genotypes of multiresistant Salmonella enterica subsp. enterica serovar Typhimurium var. Copenhagen isolates from animal sources. J Antimicrob Chemother 51:180–182.

64. Gebreyes WA, Altier C. 2002. Molecular characterization of multidrug-resistant Salmonella enterica subsp. enterica serovar Typhimurium isolates from swine. J Clin Microbiol 40:2813–2822.

